# Loose social organisation of AB strain zebrafish groups in a two-patch environment

**DOI:** 10.1101/440149

**Authors:** Axel Séguret, Bertrand Collignon, Léo Cazenille, Yohann Chemtob, José Halloy

## Abstract

We study the collective behaviour of zebrafish shoals of different numbers of individuals (1, 2, 3, 5, 7, 10 and 20 AB zebrafish *Danio rerio*) in a constraint environment composed of two identical square rooms connected by a corridor. This simple set-up is similar to a natural patchy environment. We track the positions and the identities of the fish and compute the metrics at the group and at the individual levels. First, we show that the number of fish affects the behaviour of each individual in a group, the cohesion of the groups, the preferential interactions and the transition dynamics between the two rooms. Second, during collective departures, we show that the rankings of exit correspond to the topological organisations of the fish prior to their collective departure. This spatial organisation appears in the group a few seconds before a collective departure. These results provide new evidences on the spatial organisation of the groups and the effect of the number of fish on individual and collective behaviours in a patchy environment.

## I. INTRODUCTION

Across the collective behaviours observed in social animals, collective movements [1–8], nest site selections [9–12] and site transitions [13] have been evidenced in many species. In this latter case, the groups face several alternatives and transit between them. The study of these transitions relies on decision-making processes and individual or collective preferences for environmental [14] or group members characteristics [3, 15–17] like leadership [18], motion [19] or behavioural traits, for example bold and shy individuals [12, 20].

Numerous animal species have been observed in different sorts of constraint setups or mazes to study collective movement from one site to another: corridor type [3, 16, 21], Y-maze [22], T-maze [23] or Plus-maze [24, 25]. Such constraint set-ups engage the animals to transit alone or in group from site to site and allow the observation of leadership [27–29], initiation of group movements [19, 27, 30], followers organisations [27], pre-departure behaviours [19, 28] and sites transitions [13, 31, 32]. In these latter cases the authors studied the transitions from one site to the other of one and two fish separated by a transparent partition (*Gasterosteus aculeatus* and *Sciaenops ocellatus*). Although such experimental procedure provided evidence of different leader/follower behaviours in fish, they prevent the fish from direct interactions between each other during the departures.

On the one hand, studies performed with groups of fish swimming together have evidenced that the group size can impact swimming behaviours with a variety of results. [33–35] showed that the speed, the turning speed, the nearest neighbour distances, the milling or the alignment are affected by the number of group members. The authors present opposite results depending on the species: increasing the group size of *Oreochromis niloticus* (330 and 905 fish), makes a stronger alignment [33], while for *Notemigonus crysoleucas* (30, 70, 150 and 300 fish) alignment decreases [35]. On the other hand, *Shelton et al.* [36] have shown that the density influenced nearest neighbour distances in *Danio rerio* when *Frommen et al.* [37] noticed that shoaling preferences might not always be influenced by a higher number of group members but also by the density and cohesiveness of the respective groups.

We focus on the collective movements between two environmental patches of different numbers of zebrafish. We have shown in a previous study that zebrafish transit without interruption from one landmark to another one in an open environment during trials of one hour [38]. Moreover, we have shown that groups of fish were swimming along the border of the tank and thus had a strong thigmotactic tendency [39]. Inspired by the experiments developed for highly dynamical groups of animals like the ants [40] or the fish [3, 16, 21] in constraint set-ups, we created a binary choice set-up able to channel the groups of zebrafish and to increase their stabilisation in the patches. Our experimental set-up is composed of two environmental patches (rooms) linked by a corridor (see Figure ??). The geometry of the setup is designed to study collective transitions between patches allowing to quantify the group cohesion and collective decision-making. In this study, we aim at characterising the dynamics of departure during sites transitions for several group sizes (1, 2, 3, 5, 7, 10 and 20 individuals) of AB zebrafish swimming in a constraint environment. Here we consider group size as the number of fish in a group.

Zebrafish are a gregarious vertebrate model organisms often used in behavioural studies [41, 42]. In the laboratory as much as in the nature, the zebrafish behave in groups [3, 43, 44]. They are native to the Indian sub-continent and live in small groups or in big shoals of several hundreds of fish depending on the region and the water or the environmental features (temperature, pH, human activity, predators, …) [15, 45–47]. Zebrafish live in a wide variability of habitats with varying structural complexities [46, 48] (from river channels, irrigation canals to beels) and we based our experimental method on the observations of fish swimming in a constraint set-up composed of two identical squared rooms (evoking patchy environments [49]) connected by a long corridor.

Here, we study the collective dynamics of group transitions in zebrafish with a new type of set-up. By observing groups composed of different numbers of fish, we evaluate the influence of the number of individuals in the shoals on the structure of the group (cohesiveness, inter-individual distance) and on the sequence of exit for each collective departure. By performing trials of one hour, we could observe a large number of successive transitions.

## II. RESULTS

### A. Group structure and number of individuals

First, we studied the change of the group structure according to the location of the group and the number of fish by measuring the nearest neighbour distances for each individual. Figure 1 shows the boxplots of the medians of the nearest neighbour distance distributions for each fish in 6 shoal sizes (1, 2, 3, 5, 7 and 10 fish). Thus, the boxplots for each area (rooms or corridor) and each number of individuals consist in 12 values of medians. For groups of 2 to 5 individuals, the increase of the number of individuals made the medians of the nearest neighbour distances decrease until a plateau value of approximately 4 cm. For groups of 5, 7 and 10 individuals, the medians of the nearest neighbour distances remained very close from each other.

We compared with a Two-way ANOVA the distributions of the medians of the nearest neighbour distances for each fish focusing on each area (room 1, room 2 and corridor) or each number of individuals. The test shows that there is an effect of the number of individuals on the medians of the nearest neighbour distances (*p* – *value* < 0.005, *F* = 3.87, *MS* = 0.00092 and *df* = 4). However, it does not show any significant effect of the type of the area – Room 1, Room 2 or Corridor – (*p* – *value* > 0.1, *F* = 1.96, *MS* = 0.00047 and *df* = 2) nor of an interaction between the number of individuals and the type of the area (*p* – *value* > 0.5, *F* = 0.15, *MS* = 0.00004 and *df* = 8).

### B. Oscillations and collective departures

Then, we characterised the collective behaviour of the fish. In particular, we focused our investigation on the oscillations between both rooms and the collective departure dynamics of the groups. First, we studied the repartition of the fish among the two rooms. Approximately 70% of the positions of the fish were detected in the rooms, independently of the number of individuals (Figure S23). In the Figure 2, we show that 80% of the time, less than 20% or more than 80% of the whole group is detected in the room 1. This result highlights that, as expected for a social species, the fish are not spread homogeneously in the two rooms but aggregate collectively in the patches, with only few observations of homogeneous repartition in both rooms. However, this analysis also shows that the proportion of observation with equal repartition between both rooms (40-60%) increases with the number of individuals. Thus, even if they are mainly observed together, fish in large group have a slightly higher tendency to split into subgroups. We show that the frequencies of observations for the proportions of 80 to 100% of the whole group in the room 1 are higher than 50% for all group sizes, except for 10 and 20 fish. For each trial, we defined the room 1 as the starting room where we let the fish acclimatize during 5 minutes in a transparent perspex cylinder. This may explain the observed bias of presence in favour of room 1 that may be a consequence of longer residence time at the beginning of the trials.

Then, since the fish are observed most of the time forming one group in one of the two rooms, we studied the transitions of the majority of fish between the two patches during the whole experimental time. In Figure 3, we plot the number of transitions between both rooms (see Figure S24 and Table SIII (D) of the appendix for the plot of the means of the numbers of transitions and their standard deviations in a table). First, we present the total number of transitions (All *transitions*) for all group sizes (referred as *All transitions*). Then for groups with at least two individuals, we detailed these transitions into two subcategories: *Collective transitions* (i.e. the majority of the group is detected successively one room, the corridor and the other room) and *One-by-one transitions* (i.e. the majority of the group is detected successively in one room and then in the other room, indicating that the fish did not cross the corridor together). In addition, we also quantified the *Collective U-turns* that occur when the majority of the group was detected successively in one room, in the corridor and back to the previous room.

For larger groups, the numbers of *All transitions*, *Collective transitions* and *Collective U-turns* decrease while the number of *One-by-one transitions* increases. For the transitions (collective, one-by-one and all), this tendency intensifies for bigger groups of 10 and 20 zebrafish. Also, for groups of 3 zebrafish, there are less *Collective transitions* (as well as *All transitions*) than for groups of 2, 5 and 7 zebrafish. *U-turns* remained rare and are very stable for all shoal sizes and their highest mean numbers are reached for groups of 2 and 3 zebrafish. *One-by-one transitions* are as well very rare for small groups and increase when the shoal size reaches 10 zebrafish.

For each number of fish (Figure 3), we compared with a Kruskal-Wallis test the distributions of the number of transitions (Collective, One-by-one and U-turns) and found: for 1 fish, df = 2, Chi-sq = 31.62 *p* < 0.001; for 2 fish, df = 2, Chi-sq = 30.76, *p* < 0.001; for 3 fish, df = 2, Chi-sq = 30.94, *p* < 0.001; for 5 fish, df = 2, Chi-sq = 30.41, *p* < 0.001; for 7 fish, df = 2, Chi-sq = 30.54, *p* < 0.001; for 10 fish, df = 2, Chi-sq = 19.22, *p* < 0.001 and for 20 fish, df = 2, Chi-sq = 18.36, *p* < 0.001. For each group size, we show that at least one of the distributions is significantly different from the others. The Tukey’s honest significant difference criterion shows that: all the distributions are significantly different (*p* < 0.05) except in groups of 10 individuals between *Collective U-turns* and *One-by-one transitions* and in groups of 20 individuals between *Collective U-turns* and *Collective transitions*.

As most of the transitions occur in groups, we analysed the dynamics of collective departure from the rooms with a particular emphasis to the pre-departure period. Thus, for each collective departure of the fish, defined as the whole group leaving one of the resting sites for the corridor towards the other one, we identified the ranking of exit of each fish and also their distance from the first fish leaving the room (i.e. defined as the initiator) measured at the departure timing of this initiator. Figure 4 represents the normalised contingency table of the rank of exit for all zebrafish from both rooms (without distinction) with the rank of the distances of all zebrafish to the initiator. These results correspond to 12 replicates of groups of 5 and 10 zebrafish. The initiator has a rank of exit and a rank of distances of 1. For example, in (A) the probability that the first fish to follow the initiator (rank 2) was also the closest fish of the initiator when it exited the room is 0.82. As evidenced by the darker diagonal of the contingency matrix, the rank of exit was closely related to the distance from the initiator at the beginning of the departure. Figure S25 and S26 of the appendix show a more detailed version of the Figure 4 for 3, 5, 7 and 10 individuals. In Figure 5, we plot for 3, 5, 7 and 10 zebrafish the values of the probability of equal ranking between the exit and the distances with the initiator (i.e. the diagonal of the previous plots – Figure 4) for different time-lag before the exit of the initiator. In particular, we computed the ranking of the distance from the initiator at 1 to 5 seconds before the exit of the initiator. First, these measures show that the further from the time of the initiation the lower the probability of equal ranking. This assessment is valid for every shoal sizes. Second, we see that the probability of equal ranking is often higher for the first and for the last ranked fish even few seconds before the initiation (around 2 seconds before the initiation). In Figure 6, we use the Kendall rank correlation coefficient to see if the rank of exit and the rank of distances with the initiator are dependent (close to 1) or not (close to 0) through the time. For every group sizes, we show an increase of the Kendall rank correlation coefficient when closer to the initiation. For 3 zebrafish, the time series shows that from 4 seconds before the initiation the Kendall rank correlation coefficient fully increases from 0.11 to 0.79 (at T = t = 0 s). For 5 zebrafish, it increases from 0.06 (at T = t - 4 s) to 0.75 (at T = t = 0 s), for 7 zebrafish, it increases from 0.10 to 0.70 and for 10 zebrafish, it increases from 0.08 to 0.58. These results show that for all group sizes, the closer to the initiation the higher the correlation between the rank of exit and the rank of the distances with the initiator.

## III. DISCUSSION

We studied the impact of the number of individuals in the shoals (1, 2, 3, 5, 7, 10 or 20 individuals) on the collective motion and the collective departure between two environmental patches in adult AB zebrafish.

Here we consider group size as the number of fish in a group. By changing that number of fish it may lead to size or density effects. We do not adress the question of untangling size or density effects in this study. The density is related to a need of individual space and the group size is related to a limitation in the considerations, for each individual, of the other members of the group. Our set-up reached a compromise between reducing the size of the set-up for small numbers of individuals, that would have an effect on the small group collective transitions, and increasing the size of the set-up, that would have led to tracking issues (less pixels by fish) [50]. Actually, if we resize the set-up proportionally to the density, we will modify the lengths and the widths of the rooms and the corridor. This would have an effect on the time of residency in the rooms or on the decision to cross the corridor and would introduce new variables in the experiment.

Furthermore, the Figure 1 showing that the more individuals the lower the nearest neighbour distances elucidates the previous statement. For small group, the medians of the nearest neighbour distances distributions decrease when increasing the numbers of individuals. This result could be due to an effect of the set-up where the environment is not resized in proportion of the number of the individuals. For groups of at least 5 fish, it seems that the medians of the nearest neighbour distances distributions are very similar. Here, it is possible that the threshold value of the nearest neighbour distance for the AB zebrafish has been evidenced. In this latter case, it means that the zebrafish spread more in the set-up which show that the groups are not denser when we increase their sizes. Moreover, the results of the Two-Way ANOVA evidenced that the influence of the number of individuals over the nearest neighbour distances was significant when the influence of the type of the area (rooms or corridor) is not. In [51], the authors have shown that the mean of nearest neighbor distances is about 30 mm for groups of 8 wild type zebrafish (presumably AB). Their results are very close from ours and the experiments were performed in a circular shape bowl of 20 cm in diameter. Both of our results may show that the environment has no influence on the compact structure of big groups. In parallel, if we focus on the distances between all pairs within a group we show that the higher the number of individuals the higher the medians of the distances between all respective pairs of zebrafish (Figures S11, S12, S13, S14, S15 and S16 of the appendix). Finally, we show that the higher the number of individuals the lower the time the fish stay with their nearest neighbours (Figure S 17). The combination of these results shows that there is a clear effect of the number of individuals on the cohesion; an effect that we already have shown in [38] where the bigger the group the higher the cohesion of the whole group.

It seems that there are preferential interactions between zebrafish (Figure S11, S12 and S13 of the appendix) and increasing the number of individuals will affect these interactions: respective pairs are less cohesive in larger groups. We considere that revealing the time spent by the fish with their nearest neighbours show stronger informations about preferences between pairs than the median distances of each couple, which are already approximations. Hence, we show (Figure S 17) that for goups of 3 fish the distribution of the number of times the fish stay with their nearest neighbours is not significantly different from a random uniform distribution. It means that for such number of individuals there is no preferential interaction. For bigger groups, these distributions are significantly different from a random uniform distribution which means that there are preferential interactions. Preferential interactions have been evidenced in other species: Briard et al. [59] show affinities, hierarchy and pairs interactions in a group of domestic horses, [60–63] show that the affinity between individuals (monkeys, *Macaca mulatta, Macaca tonkeana, Papio ursinus*; or fish, *Gasterosteus aculeatus*) play a role in the collective movements. We propose two hypotheses that could explain the change of the interactions between pairs of zebrafish when changing the number of individuals. On the one hand, in groups larger than two fish, each zebrafish has to choose the preferred partners, between all other fish. In larger groups there are more individual choices and more preference tests. On the other hand, the patchy environment may break pair interactions and may force the emergence of new pairs. These two hypotheses could explain the dynamics of the pair interactions observed during the experiments.

The fish are detected 70% of the time in the rooms (Figure S 23). On average, they spend about 10 seconds in a room (Figure S 22), transit to the other room through the corridor (4 seconds on average) and then come back. They oscillate between the rooms. In a previous study we showed that zebrafish also transit and oscillate between landmarks in an open environment [38]. The Figure 3 shows that most of the transitions are collective when the Figure 2 shows that the whole group swim generally together in both rooms. This observation is strengthened by the very rare number of *One-by-one* transitions between the rooms. However, groups of 10 and 20 zebrafish show sharp decreases in the number of collective transitions. This drop could be due to the topology of the set-up and congestion effects. Larger groups can split into smaller subgroups. The threshold we imposed in the analysis of the collective transitions (below 70% of the whole group, the transitions were not taken into account) may reinforce this effect. This seems to be confirmed by the Figure S 24 which shows the mean and median number of transitions for different numbers of individuals when all the fish start to move from a room: the larger the group, the lower the number of transitions with the whole group. Also the Figure 2 shows that the bigger the groups, the higher the frequency of observations of sub-groups located in the rooms and in the corridor (20-40%, 40-60% and 60-80%). Many studies have analysed the fusion-fission mechanisms occurring in groups of fish or mammalians. [64–66] show that these mechanisms are frequent in the wild and generate body length assortment within groups of fish (*Fundulus diaphanus, Notemigonus crysoleucas, Catostomus commersonii, Poecilia reticulata*). Sueur et al. [8] show that fission-fusion mechanisms participate in the information transfer between subgroups and the group of *Myotis bechsteinii*.

In the corridor, we observe few u-turns. The zebrafish swimming preferentially along the walls and a canalisation effect of the corridor may explain this observation. As expected, connecting the two patches, the corridor is used as a mere transit area (zebrafish show higher speeds in the corridor – Figure S 19).

We show that the organisation of the group during collective departure takes place during a short pre-departure period and is related to the distances between the initiator (of the exit from the room) and the other fish. Ward *et al.* has shown that the first fish (of a group of 5 *Dascyllus aruanus*) to follow the initiator is generally (rank = 2: 53% of the trials over 2 trials for each 15 groups of fish) the nearest neighbour of the initiator and that the frequency of equality between the rank of exit and the rank of the distances from the initiator decreases, with these results rank = 3: 27% and 33%, rank = 4: 20% and 7% then rank = 5: 0% and 7%) [27]. We tested four different numbers of individuals (3, 5, 7 and 10 zebrafish) and show similar results especially on the decreasing trend of the probability when focusing on the next ranked fish (Figures S 26, S 25 of the appendix and Figure 5). However, we observe an extremely high probability of equal ranking for the first fish that follows the initiator (rank = 2: 75% to 90%), high probabilities of equal ranking for the second and the last fish that follow the initiator (rank = 3: 50% to 65% and rank = last fish: 38% to 75%) and show that the probabilities of equal ranking for the other fish are quite similar to each others. The rank of exit and the rank of the distances from the initiator are strongly correlated at the moment of the exit (T = 0s, Figure 6), from 60% to 80%. Hence, it seems that the organisation of the zebrafish groups (2^*nd*^, 3^*rd*^ and last ranked fish) during the collective departures is topological. Other studies about the organisations of collective departures show a joining process for *Equus ferus caballus* that is related to affinities and hierarchical rank [59]. Rosenthal *et al.* show that, in groups of *Notemigonus crysoleucas*, the initiator is the closest fish from the group boundary in 27% of the cases and the first responder is the closest fish from the group boundary in 19% of the cases [30]. Moreover during the initiation, when fish leave the rooms, our results suggest the idea of cascades of behavioural changes already developed by Rosenthal *et al.* [30]: the initiator drags another fish along that drags another one, etc.

This organisation appears a few seconds before the fish leave a room to transit to the other one. Two seconds before the initiation, the group show a structure that prepares for the exit (Figure 5). The Kendall rank correlation coefficient confirmed the idea of the organisation as it reaches 18% to 25% two seconds before the departure and 30% to 50% one second before the departure (Figure 6). In the literature we found other cases of initiations: [20, 67] have shown that a three-spined stickleback *Gasterosteus aculeatus* or a sheep *Ovis aries* alone moving away from the herd can initiate a collective departure, [68] have notice a large variety of intiations for groups of mountain baboons *Papio ursinus* where the initiator can be joined by the group immediately or [69, 70] have observed for white-headed capuchins *Cebus capucinus* a synchronization of their behaviours and a minimum proportion of the whole group is able to launch a collective departure.

In conclusion, this study shows that the number of fish affects the motion of each individual in the groups and the group cohesion. The analysis of the dynamics shows that the zebrafish oscillate mainly in groups between the two patches in the environment and that the majority of the departures is collective. During the collective departures, we observe that an intra-group organisation appears prior to the transition. Increasing the number of individuals makes this organisation less predictable. Finally, we noticed that a few seconds before the collective departures the groups have a particular spatial organisation.

## IV. METHODS

### *1.* Fish and housing

We bred 600 AB strain laboratory wild-type zebrafish (*Danio rerio*) up to the adult stage and raised them under the same conditions in tanks of 3.5L by groups of 20 fish in a zebrafish aquatic housing system (ZebTEC rack from Tecniplast) that controls water quality and renew 10% of water in the system every hour. Zebrafish descended from AB zebrafish from different research institutes in Paris (Institut Curie and Institut du Cerveau et de la Moelle Épinière). AB zebrafish show zebra skin patterns and have short tail and fins. They measured in mean = 3.0 cm 0.36 cm, median = 2.9 cm long. All zebrafish used during the experiments were adults from 7 to 8 months of age. We kept fish under laboratory conditions: 27 °C, 500*μ*S salinity with a 10:14 day:night light cycle, pH is maintained at 7.5 and nitrites (NO^2−^) are below 0.3 mg/L. Zebrafish are fed two times a day (Special Diets Services SDS-400 Scientic Fish Food).

### *2.* Experimental setup

The experimental tank consisted in a 1.2 m × 1.2 m tank confined in a 2 m ×2 m × 2.35 m experimental area surrounded by white sheets, in order to isolate the experiments and homogenise luminosity. A white opaque perspex frame (1 m × 1 m × 0.15 m - interior measures) is placed in the center of the tank. This frame helped us to position the two rooms and the corridor. The squared rooms (0.3 m × 0.3 m) and the corridor (0.57 m × 0.1 m) have been designed on Computer-Aided Design (CAD) software and cut out from Poly(methyl methacrylate) (PMMA) plates of 0.003 m thickness. Each wall are titled, (20° from the vertical) to the outside with a vertical height of 0.14 m, to avoid the presence of blind zones for the camera placed at the vertical of the tank. The water column had a height of 6 cm, the water pH is maintained at 7.5 and Nitrites (NO^2−^) are below 0.3 mg/L. The experiments are recorded by a high resolution camera (2048 px × 2048 px, Basler Scout acA2040-25gm) placed above the experimental tank and recording at 15 fps (frame per second). The luminosity is ensured by 4 LED lamps of 33W (LED LP-500U, colour temperature: 5500 K - 6000 K) placed on each corner of the tank, above the aquarium and directed towards the walls to provide indirect lightning.

### *3.* Experimental procedure

We recorded the behaviour of zebrafish swimming in the setup during one hour and did 12 replicates with groups of 1, 2, 3, 5, 7, 10 and 20 zebrafish. Every six replicates the setup is rotated by 90 ° to prevent potential environmental bias (noise, light, vibrations, …). Before each replicate, the starting chamber, from which the fish are released, is chosen randomly. We called the starting chamber *Room 1.* Then, the fish are placed with a hand net in a cylindrical arena (20 cm diameter) made of Plexiglas in the centre the selected rooms. Following a five minutes acclimatisation period, this cylinder is removed and the fish are free to swim in the setup. The fish are randomly selected regardless of their sex and each fish is never tested twice to prevent any form of learning. The water of the tank is changed every week and the tank and hthe set-up are cleaned during the process.

### *4.* Tracking & data analysis

Today, many studies on animal collective behaviours use methodologies based on massive data gathering, for exemple for flies (*Drosophila melanogaster*) [72, 73], birds (*Sturnus vulgaris*) [74–76], fish (*Notemigonus crysoleucas*) [77]. Our experiments are tracked in real-time (“on-line”) by a custom made tracking system based on blob detection. Each replicate except experiments with 20 zebrafish is also tracked by post-processing (“off-line”) with the idTracker software to identify each fish and their positions [50]. Each replicate consisted of 54000 positions (for one zebrafish) to 1080000 positions (for 20 zebrafish). The idTracker software is not used for groups of 20 fish due to higher number of errors and too long computing time. For example, for a one hour video with 2 fish idTracker gives the results after 6 hours of processing and for a one hour video with 10 fish it lasts a week to do the tracking (with a Dell Percision T5600, Processor: Two Intel Xeon Processor E5-2630 (Six Core, 2.30GHz Turbo, 15MB, 7.2 GT/s), Memory: 32GB (4×8GB) 1600MHz DDR3 ECC RDIMM).

Since idTracker solved collisions with accuracy [50] we calculated individual measures and characterised the aggregation level of the group (except for groups of 20 individuals). We also calculated the distances between each pair of zebrafish respectively, the travelled distances of each individual and their speeds. The calculation of the speed has been done with a step of a third of a second in sort of preventing the bias due to the tracking efficiency of idTracker that does not reach 100% (see Table SII of the appendix). The data gathered for groups of 20 individuals are only used in the analyses which focused on group behaviour and do not need the identities of the fish.

The Figure 2 has been obtained following this process: At each time step with at least one fish detected in a room, we analysed the repartition of the group among the rooms by computing the proportion of fish present in room 1=*R*_1_ / (*R*_1_+*R*_2_) with *R*_1_ and *R*_2_ the number of fish in the respective room number.

When all fish were present in the same room, we identified which fish initiates the exit from the room, established a ranking of exit for all the fish and calculated the distances between all zebrafish to the initiator to establish a ranking of distances. Finally, we confronted these ranks and count the number of occurences for each ranking case. We checked the results for different time steps before the initiation. The idea was to highlight a correlation between the spatial sorting and the ranks of exit and also a possible prediction of ranking of exit.

We looked at majority events defined as the presence of more than 70% of the zebrafish in one of the three areas of the setup, either in the room 1 or in the room 2 or in the corridor. To compute their numbers, we averaged the number of fish over the 15 frames of every second. This operation garanties that a majority event is ended by the departure of a fish and not by an error of detection during one frame by the tracking system. We then computed the durations of each of those events and counted the transitions from a room to the other one and sort them. All scripts were coded in Python using scientific and statistic libraries (numpy, pylab, scilab and matplotlib).

Finally, we computed the neighbour distances as the distances between each nearest fish (figure S 20).

### *5.* Statistics

All scripts were coded in Matlab and Python using statistic libraries (numpy, pylab, scilab and matplotlib). For the Figure S 19 we report the number of values of the speeds on the Table SI of the appendix. For the Figures 1 and 3 we tested the distributions using Kruskal-Wallis tests completed by a post-hoc test: Tukey’s honest significant difference criterion. For the Figure S 19 we tested with ANOVA N-way 10 samples of 0.5% of the data of the speed choosen randomly for each number of individuals.

The Kendall rank correlation coefficient [78], τ, is a measure of ordinal association between two measured quantities. It goes to 0 when the two quantities are independent and to 1 if they are correlated. It is computed by:

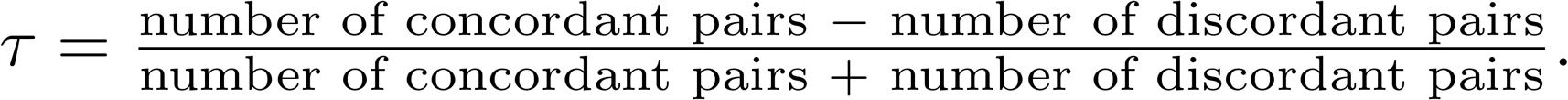

We use the Kendall rank correlation coefficient to see if the rank of exit and the rank of distances with the initiator are dependant or not through the time.

## Animal ethics

The experiments performed in this study were conducted under the authorisation of the Buffon Ethical Committee (registered to the French National Ethical Committee for Animal Experiments #40) after submission to the state ethical board for animal experiments.

## Data Availability

Supporting are available on figshare: https://doi.org/10.6084/m9.figshare.5151247.v1

## Authors’ contributions

AS carried out the lab work, the data analysis and the design of the set-up and drafted the manuscript; BC carried out the statistical analyses and participated in the data analyses and in the writing of the manuscript; LC developed the online tracker, stabilized the video aquisition and improved the idTracker; YC carried out the lab work, the design of the set-up and the data analysis; JH conceived the study, designed the study, coordinated the study and helped draft the manuscript. All authors gave final approval for publication.

## Acknowledgments

The authors thank Filippo Del Bene (Institut Curie, Paris, France) and Claire Wyart (Institut du Cerveau et de la Moelle Épinière, Paris, France) who provided us the parents of the fish observed in the experiments reported in this paper.

## V. SUPPLEMENTARY INFORMATION AND FIGURES

Supplementary figures of “Loose social organisation of AB strain zebrafish groups in a two patches environment”.

### A. Group structure and group size

We showed that the distances travelled by the zebrafish are related to the size of the group (Figure S 18 of the appendix). Groups of 2 to 7 zebrafish travelled the longer distances (with a declining trend) and fish alone and groups of 10 zebrafish travelled the shorter distances.

Figure S 19 shows the means and the medians of the individual speeds of the fish measured during the entire experimental time (one hour) and according to their spatial location (in the corridor or in one of the two rooms). The fastest individuals are observed in groups of 5 fish in the corridor and 3 fish in both rooms. On the contrary, fish alone and groups of 10 individuals show the slowest mean and median speeds. Moreover in the corridor, between the group sizes of 1 and 5 individuals, there is an increase of the mean and median speed. Then, for bigger group sizes, the means and medians decrease. Likewise, in both rooms, for the group sizes of 1 and 3, we observe an increase of the mean and the median of the speeds, then for bigger group sizes a drop.

We tested with ANOVA N-way (which is a generalisation of the ANOVA Two-way and because of the unbalanced size of the samples) 10 samples of 0.5% of the data of the speed randomly choosen for each group size. For the location (room 1, room 2 and corridor), df = 2, the F-value oscillates between 272.98 and 350.67, for the group size, df = 5, the F-value oscillates between 45.86 and 90.46, for the interaction, df = 10 and F-value oscillates between 2.07 and 4.56. There is an effect of the size of the groups and another one of the area (rooms or corridor) on the speed (p-value < 0.005). There is a small effect of the interaction of the size of the groups and the area on the speed (p-value < 0.02).

We show that the individual speed of the zebrafish varies according to the areas in which they are swimming and their group size. The zebrafish move faster in the corridor and have similar lower speeds in both rooms (Figure S 19 of the appendix). The surface of the corridor is the third of a room and it constraints the direction the fish have to follow. We have shown that zebrafish are known to swim along the walls of the experimental tank [38, 39] thus showing a strong thigmotaxis. In the corridor, that can be compared to a tunnel, canalised by the walls of the corridor, the zebrafish increase their individual speeds to make the transit from one room to the other. In both rooms the means and the medians of the individual speeds are at their highest levels for groups of 3 zebrafish and the maximum of the means and the medians of the individual speeds is reached for group size of 5 fish in the corridor. In parallel, in each area these means and medians are at their lowest levels for the smallest and the biggest group sizes: 1 and 10 zebrafish. Hence, in both rooms and in the corridor respectively, we have seen that from 1 to 3 individuals and from 1 to 5 individuals the individual speeds increase, when from 3 to 10 individuals and from 5 to 10 individuals, the individual speeds decrease.

First, our results confirm that the behaviour of a zebrafish alone differs significantly from the behaviour of zebrafish in groups. Isolated zebrafish travel a shorter distance and at a lower speed than zebrafish in groups. This can be the result of the stress generated by being isolated in a new environment. The stress level has been studied and [58] shows that some anxiolytics (fluoxetine and ethanol, which reduce stress level) will increase the speed and/or the travelled distances of zebrafish alone in a tank. Second, zebrafish swim faster in smaller group sizes and their speeds decrease for bigger group sizes. We observe the same trend for the travelled distances. These results may suggest a congestion effect where obstruction can affect their individual speeds and hence their travelled distances during the experimentation time [53]. Such effect has already been reported for example in the ant species *Atta cephalotes*: crowded conditions on the trail network make the velocity decrease [52]. Herbert-Read et al. present another explanation for the changes of the motions where each fish (*Gambusia holbrooki*) conforms to the group behaviour through the interaction rules between the individuals and the decisions of each individual to follow or copy their neighbour movements [34]. Although this case seems to be extreme, [54–56] have shown that fish from different species (*Perca fluviatilis, Gasterosteus aculeatus*) and *Gambusia holbrooki* can maintain particular individual behavioural traits in a social context. These changes in behaviours are found in other animal species such as birds (*Erythrura gouldiae*) that adjust their behaviour according to the personality of their partners [57].

**TABLE I:** Number of values of speeds. This table is related to Figure S?? of the article.

| number of values | Room 1 | Room 2 | Corridor |
| --- | --- | --- | --- |
| 1 fish | 54512 | 39829 | 24000 |
| 2 fish | 62941 | 61202 | 45698 |
| 3 fish | 129950 | 80216 | 73852 |
| 5 fish | 152880 | 119927 | 102647 |
| 7 fish | 253389 | 164149 | 170728 |
| 10 fish | 280719 | 218863 | 186999 |

### B. Oscillations and collective departures

To analyse the dynamics of the space occupancy in the set-up we computed the mean number of majority events and the mean durations and cumulative durations of occupancy by a majority of individuals within the three areas when a majority of the whole group is reached (Figure S 21 and Figure S 22). We define the majority as 70% of the individuals being present in the considered section of the set-up. On the Figure S 21, we find more majority events in the corridor than in room 1 or room 2 except with groups of 20 zebrafish. Whatever the size of the group, we find almost the same number of majority events inside the rooms 1 and 2. Also, in all areas we see that for groups of 10 and 20 zebrafish the bigger the group the lower the number of majority events. The difference between the number of majority events in the corridor and in both rooms is relatively stable for groups of 1, 2, 3, 5 and 7 zebrafish but decreases when increasing the size of the groups (10 to 20 zebrafish). The mean number of majority events finally reaches almost the same value when 20 zebrafish are tested in the setup (room1: 51,2; room 2: 43.5; corridor: 41.7). Table SIII of the appendix (A) shows, for the 12 replicates of each group size, the standard deviations linked with the number of majority events (related to Figure S 21). On the Figure S 22, we see that the means of the durations of the majority in each area follow a similar trend in both rooms and are longer than in the corridor. Increasing the size of the group has almost no effect on the durations in the corridor when it has an impact in the rooms, where fish stay longer in majority if the group size increases. However, for 20 zebrafish, durations decrease in all areas. Table SIII (B) of the appendix shows the standard deviations related to the Figure S 22.

**TABLE II:**
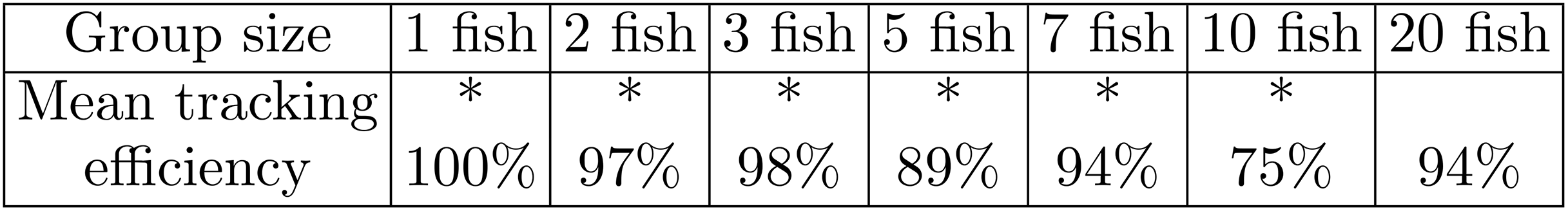
Mean tracking efficiency. * means that the experiments are tracked by the idTracker program and that we have the individual identities and the positions of the fish [50].

**TABLE III:** Standard deviations of the means of (A) majority events, (B) durations, (C) numbers of transition types with a majority of zebrafish in the three sections of the setup.

|  |  |  |  |  |  |  |  |
| --- | --- | --- | --- | --- | --- | --- | --- |
| (A) Majority events | 1 fish | 2 fish | 3 fish | 5 fish | 7 fish | 10 fish | 20 fish |
| Std. corridor | 62.5 | 42.4 | 35.7 | 41.3 | 33.3 | 49.4 | 19.2 |
| Std. room 1 | 30.2 | 17.4 | 18.8 | 21.0 | 14.5 | 26.1 | 14.3 |
| Std. room 2 | 33.0 | 28.0 | 19.8 | 21.3 | 19.8 | 24.6 | 15.8 |
| (B) Durations (s) | 1 fish | 2 fish | 3 fish | 5 fish | 7 fish | 10 fish | 20 fish |
| Std. corridor | 1.73 | 1.94 | 2.09 | 1.76 | 2.48 | 2.92 | 1.92 |
| Std. room 1 | 21.7 | 10.2 | 11.8 | 10.9 | 12.2 | 13.2 | 8.28 |
| Std. room 2 | 13.2 | 7.03 | 8.73 | 8.63 | 8.68 | 10.7 | 6.57 |
| (C) Transition types | 1 fish | 2 fish | 3 fish | 5 fish | 7 fish | 10 fish | 20 fish |
| Collective | 32.9 | 23.1 | 16.4 | 19.5 | 14.0 | 22.4 | 6.9 |
| One-by-one | 0.1 | 2.4 | 3.7 | 3.3 | 3.7 | 7.8 | 8.8 |
| U-turns | 12.0 | 12.3 | 7.0 | 6.9 | 5.3 | 5.4 | 4.5 |

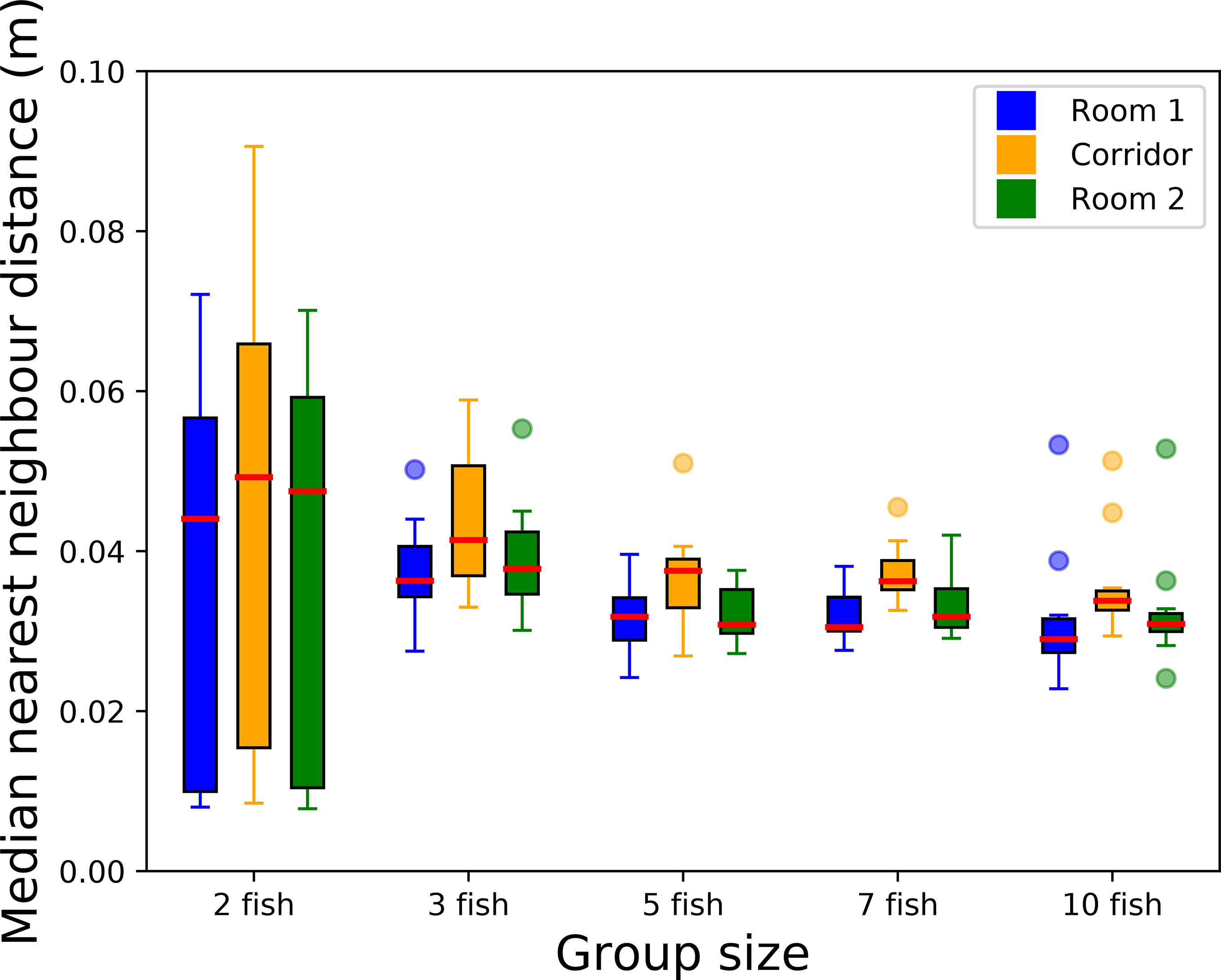

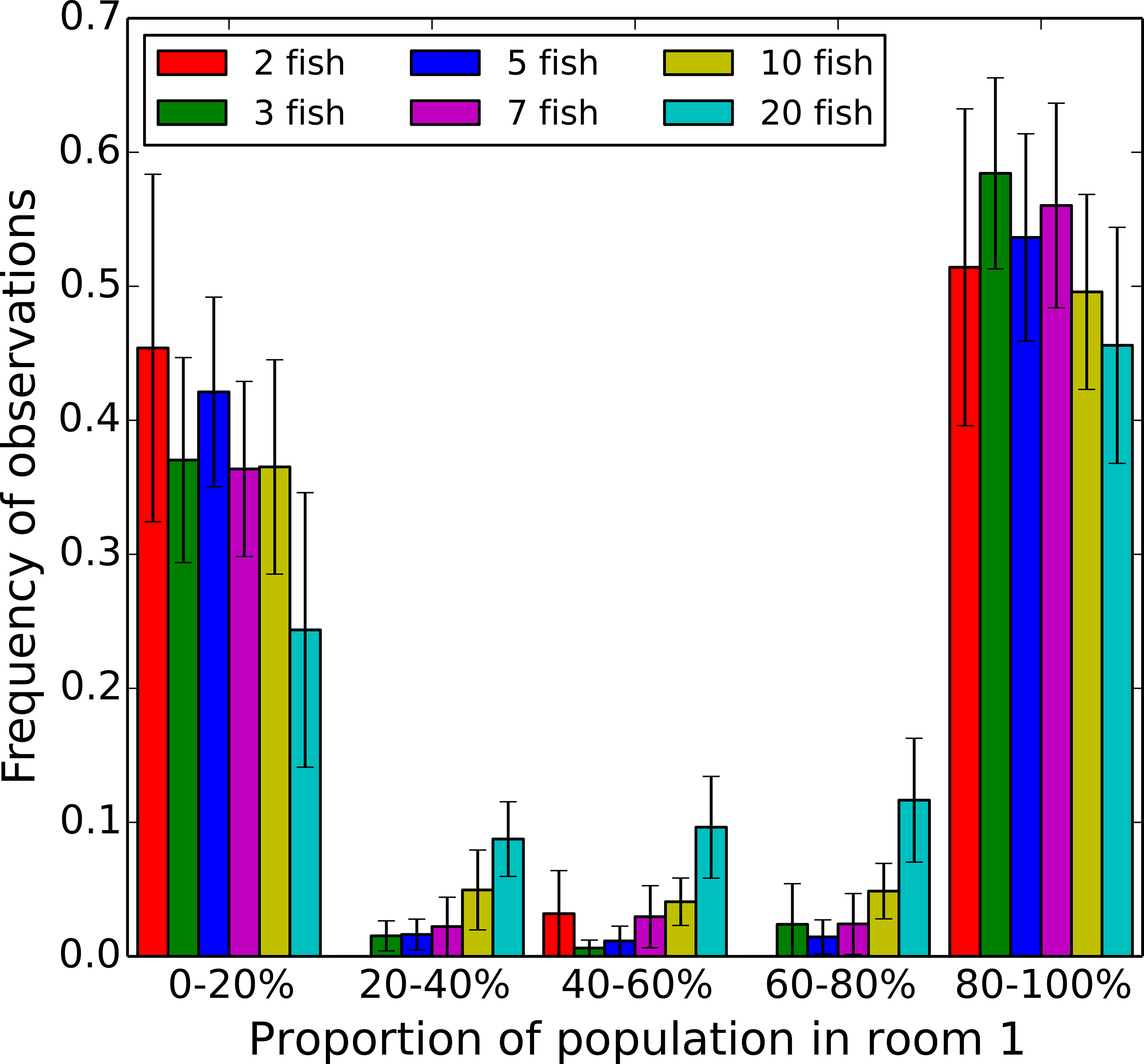

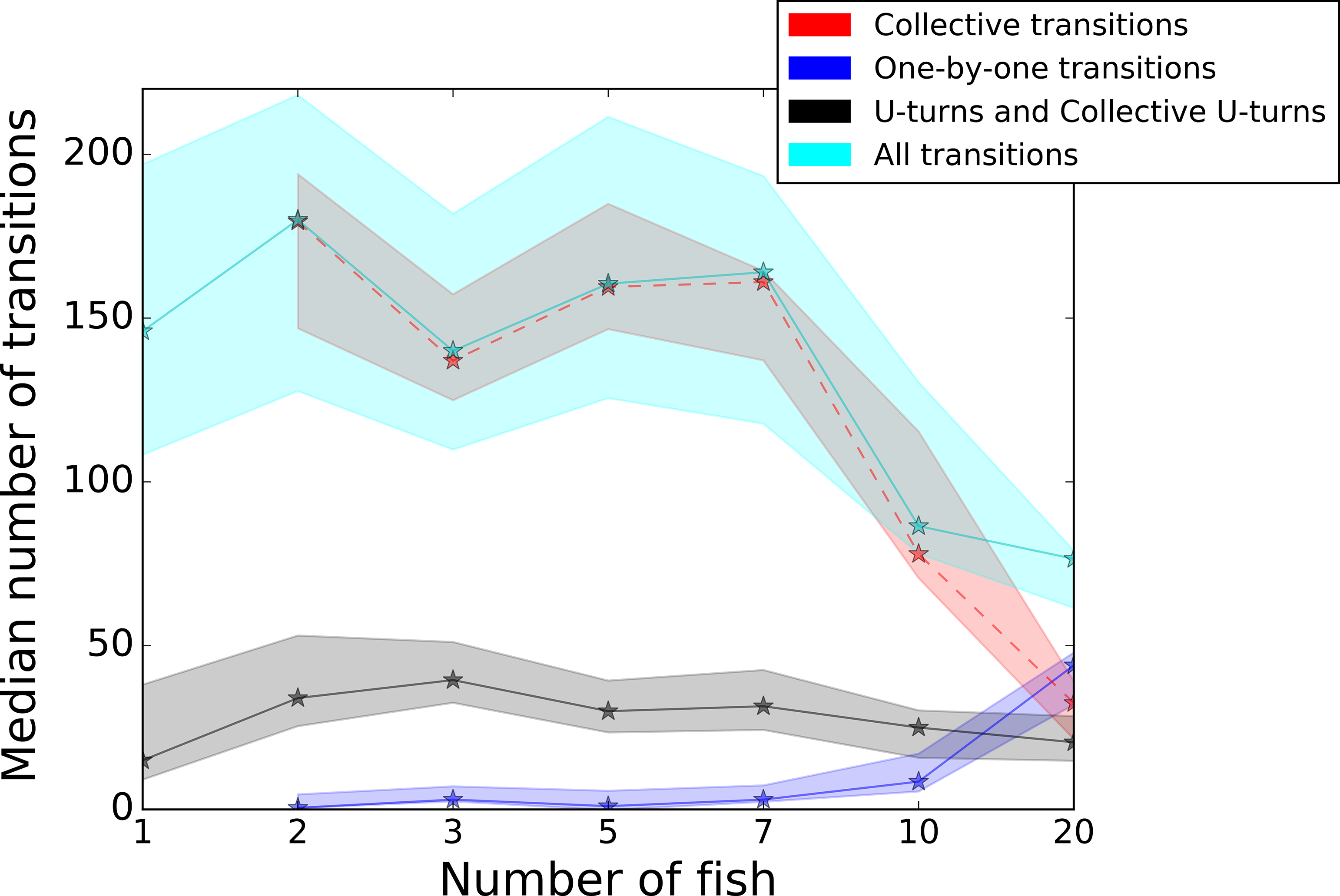

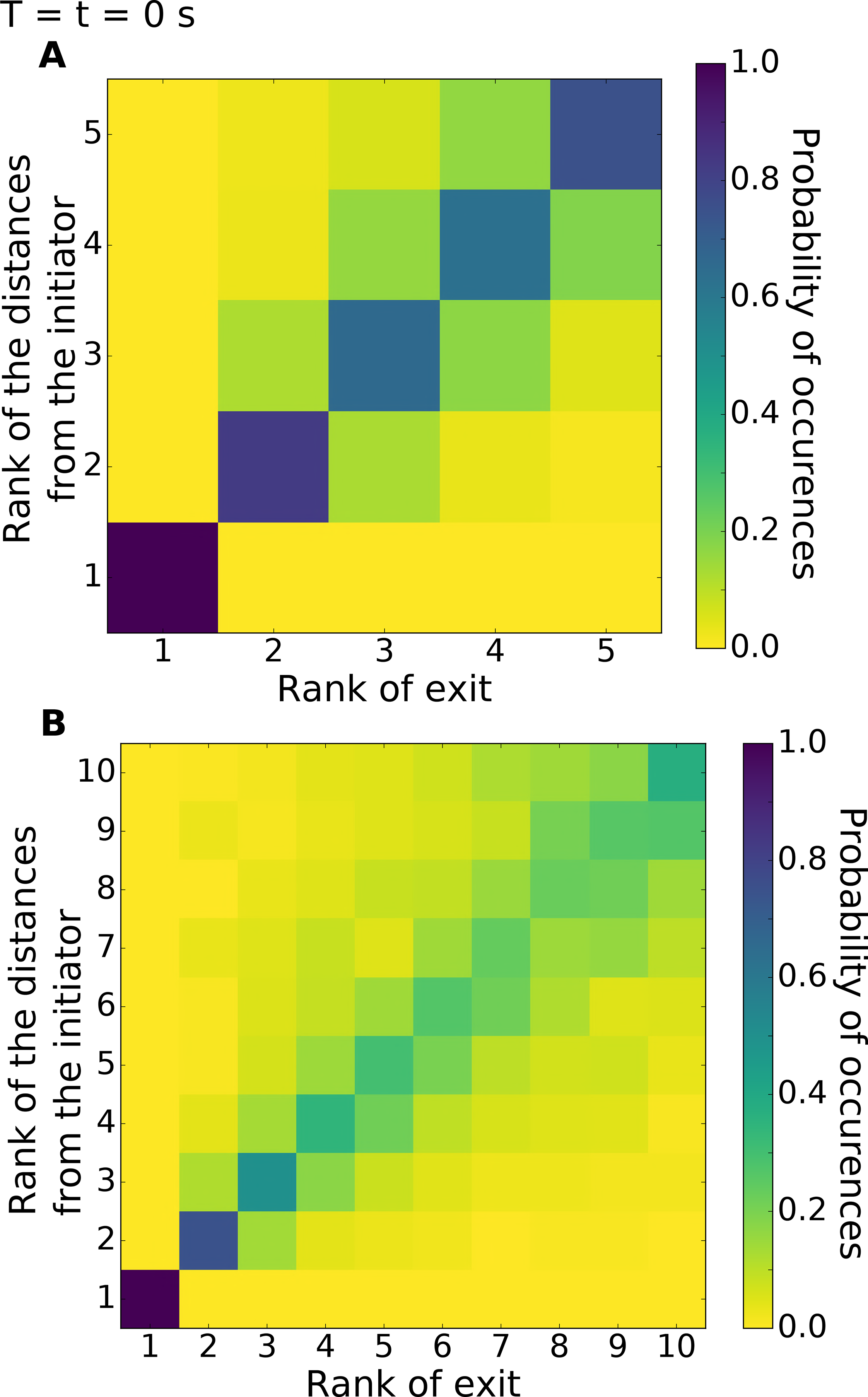

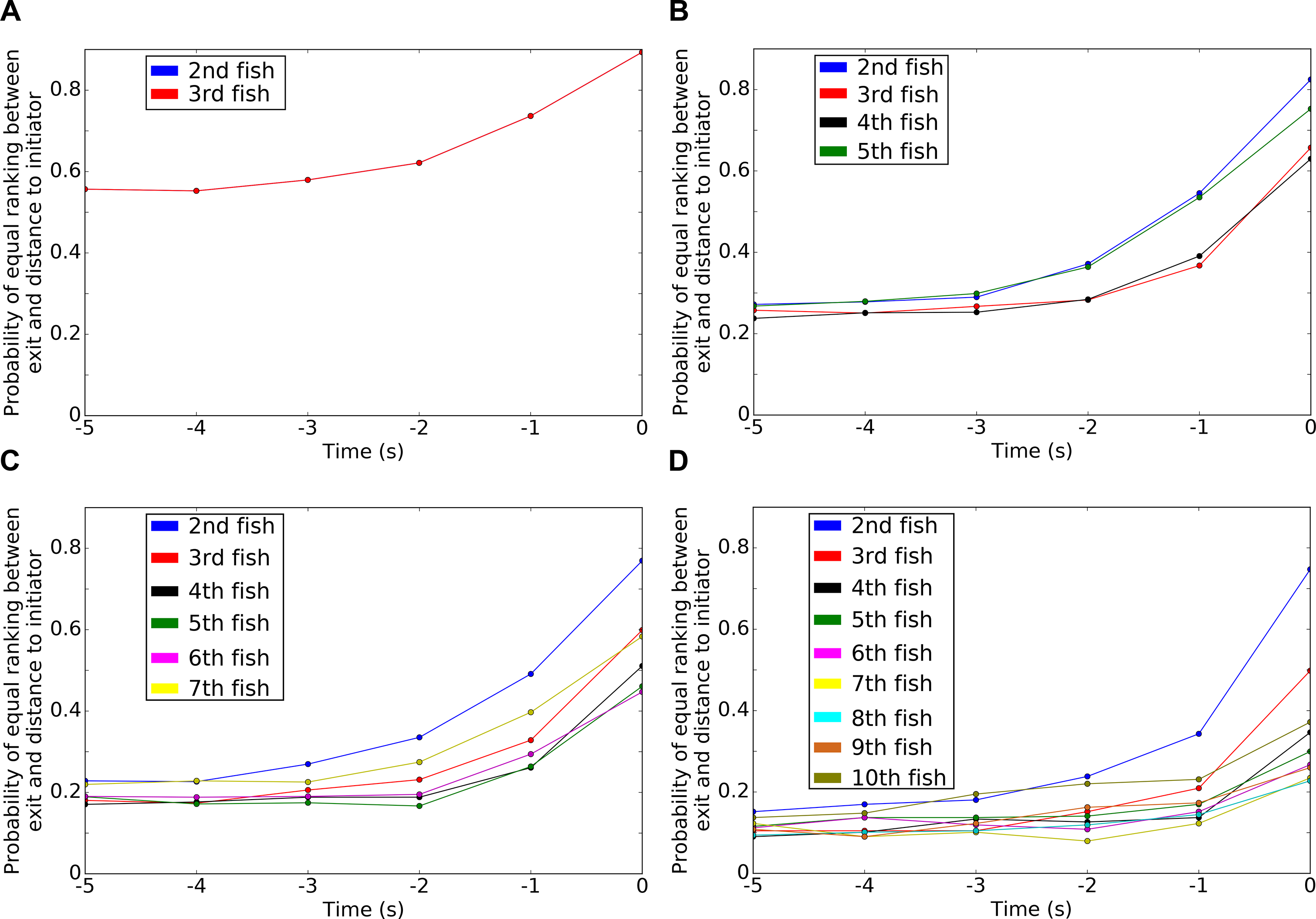

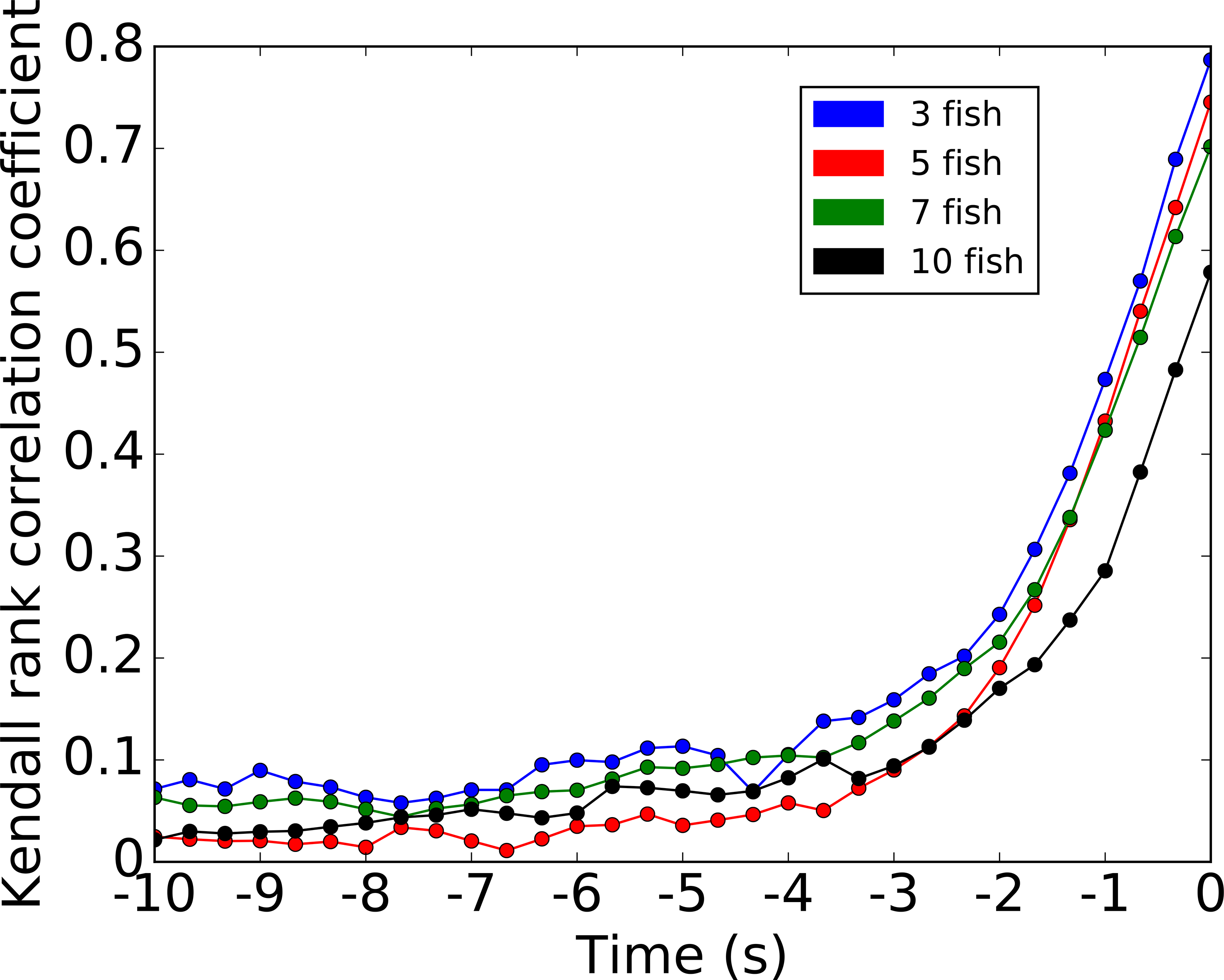

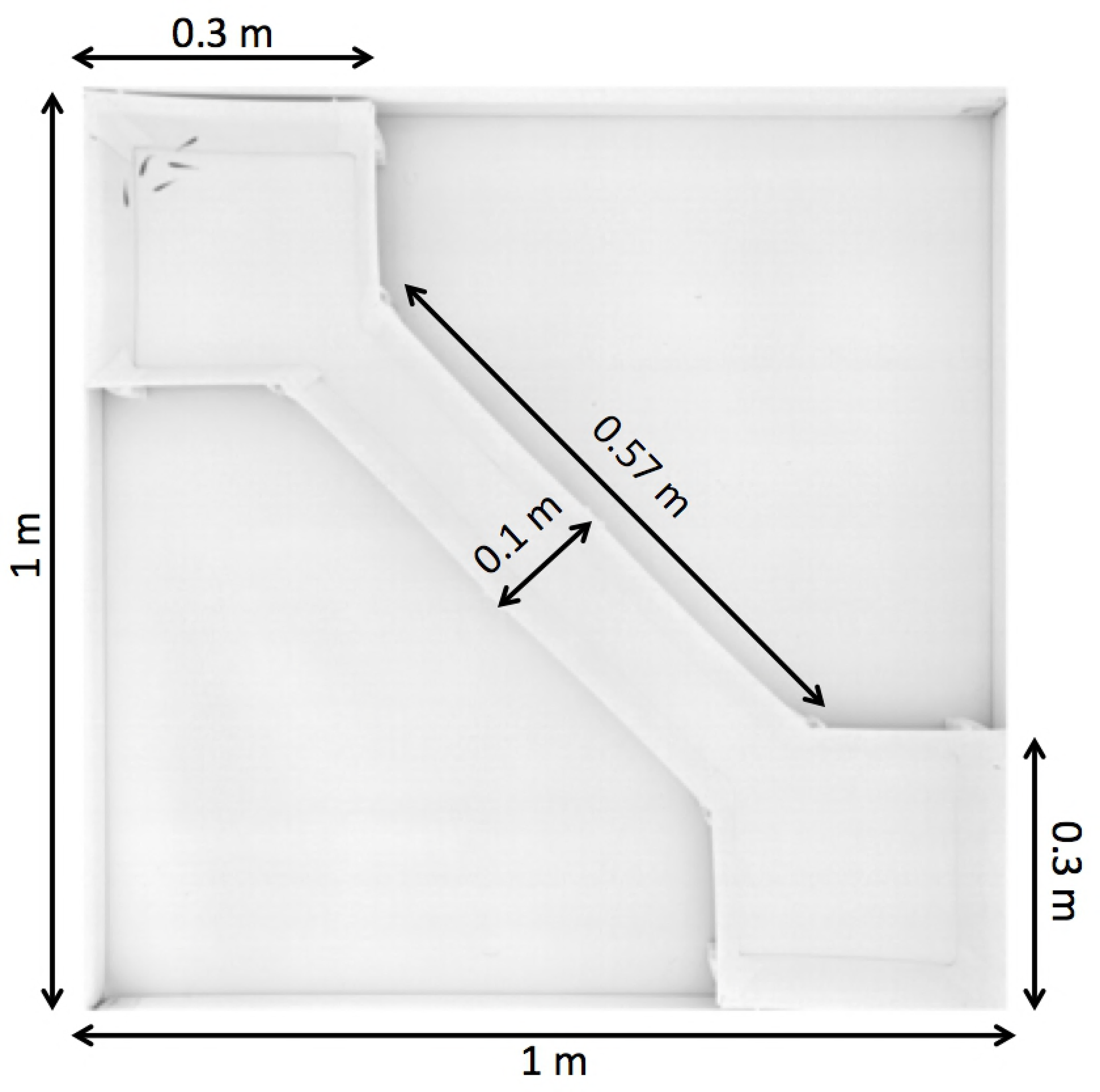

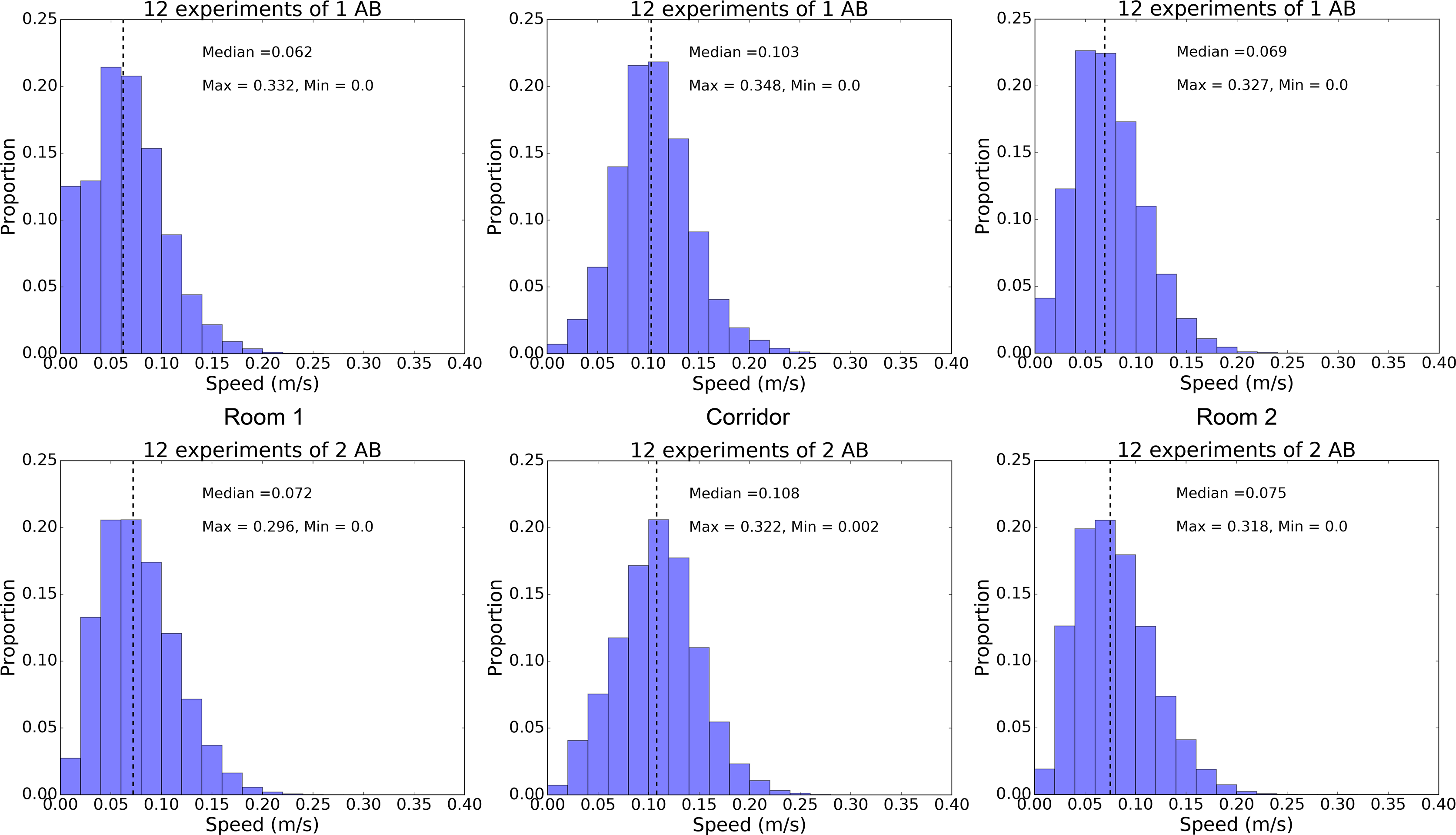

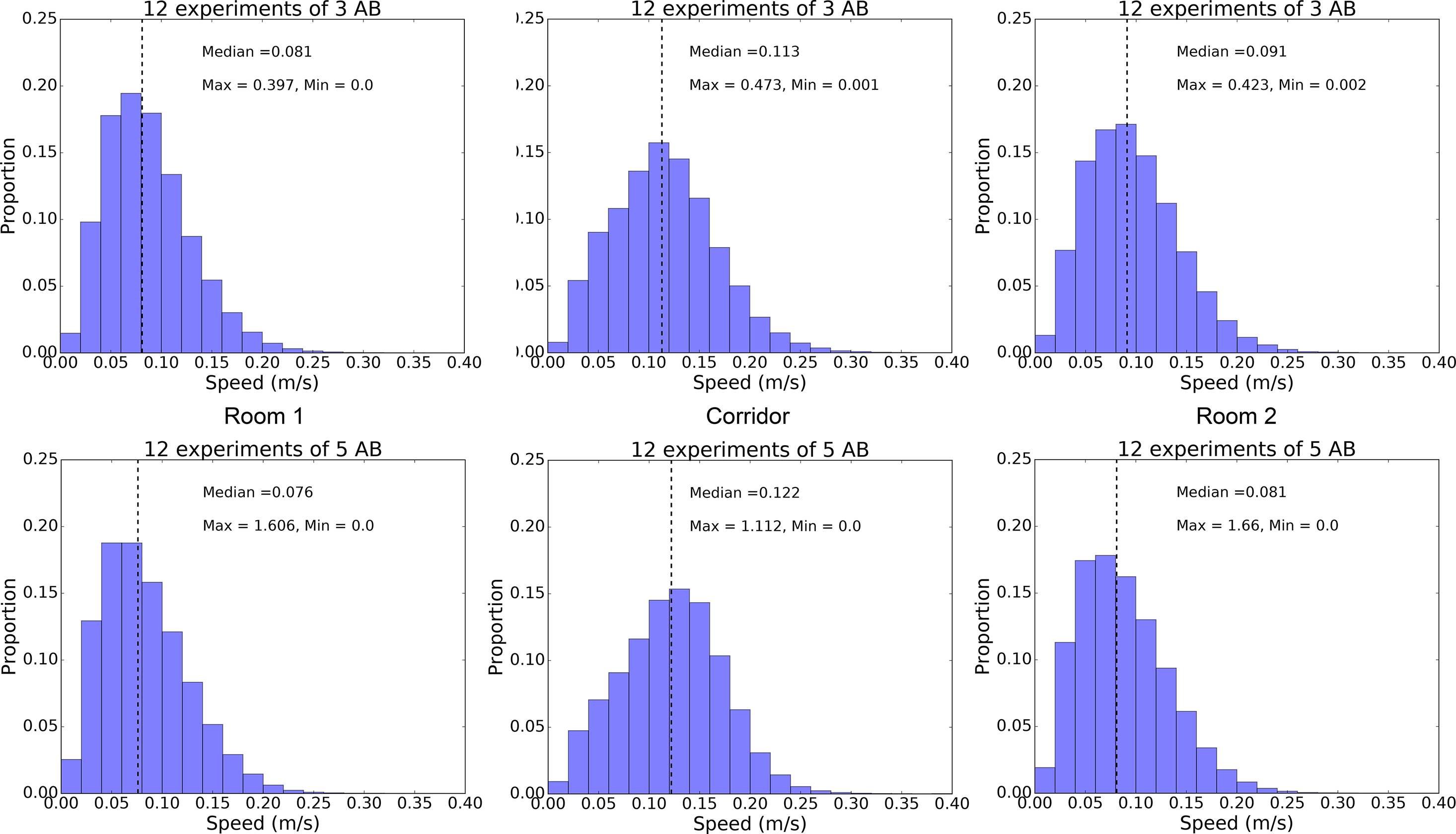

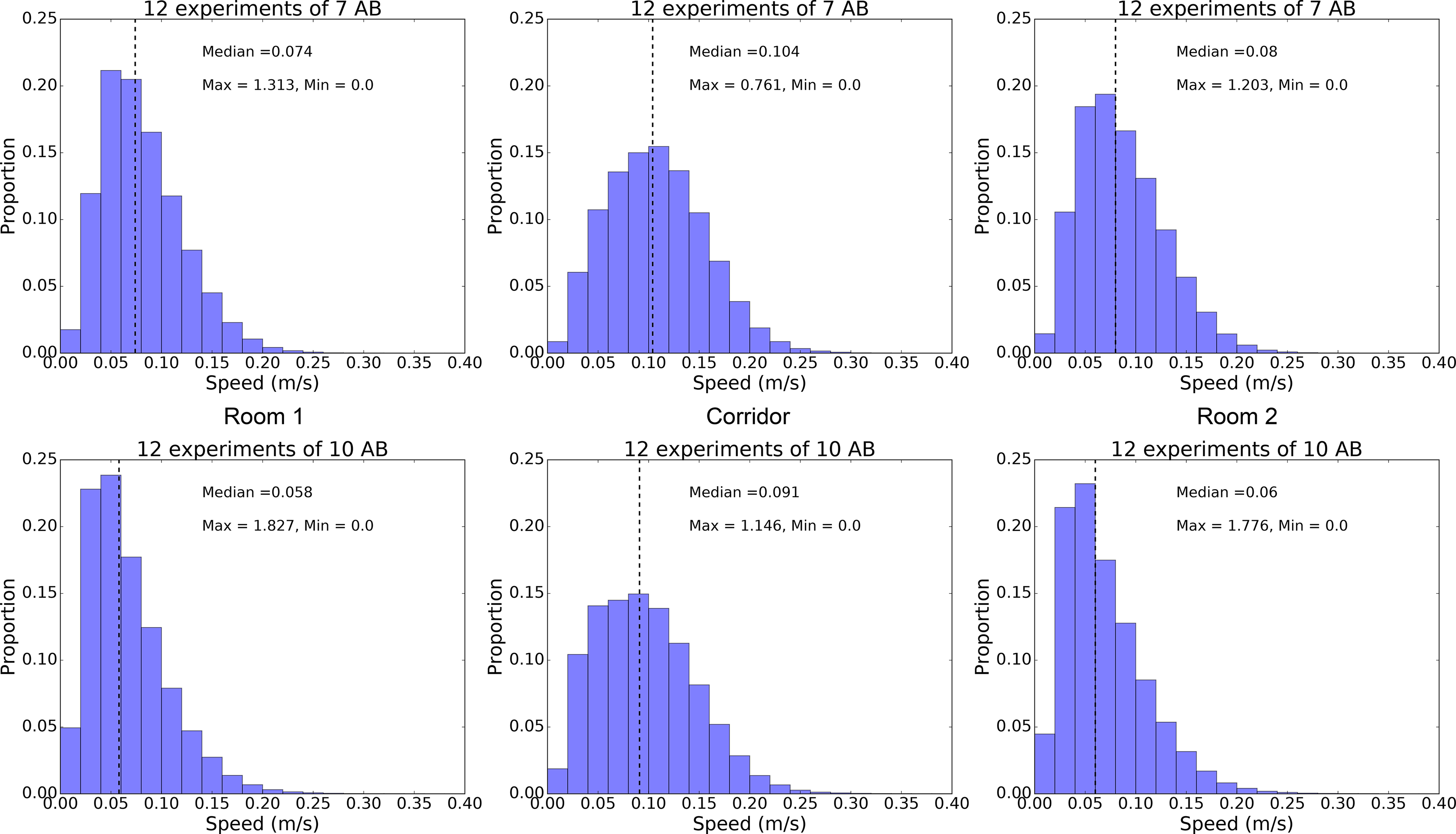

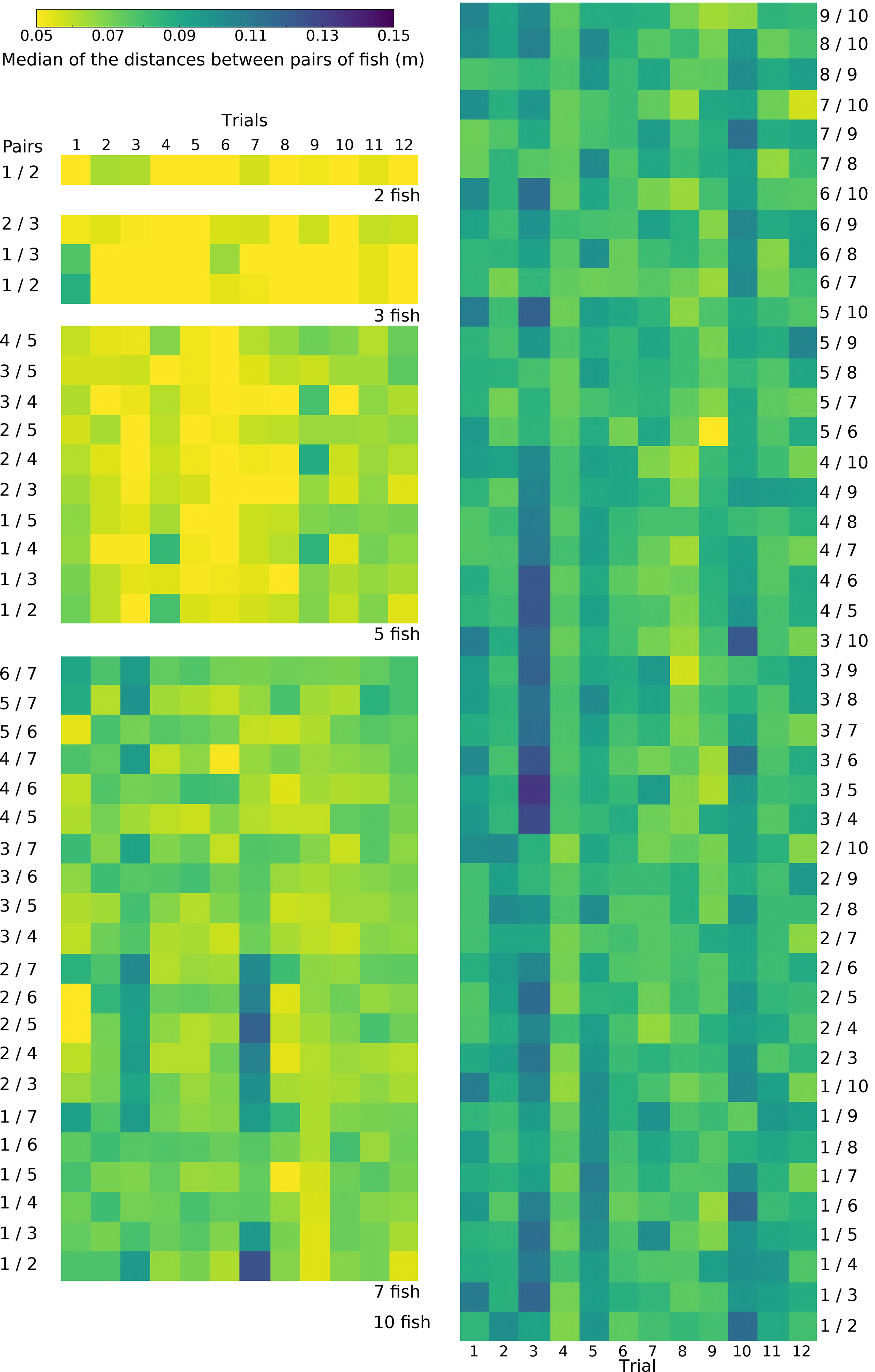

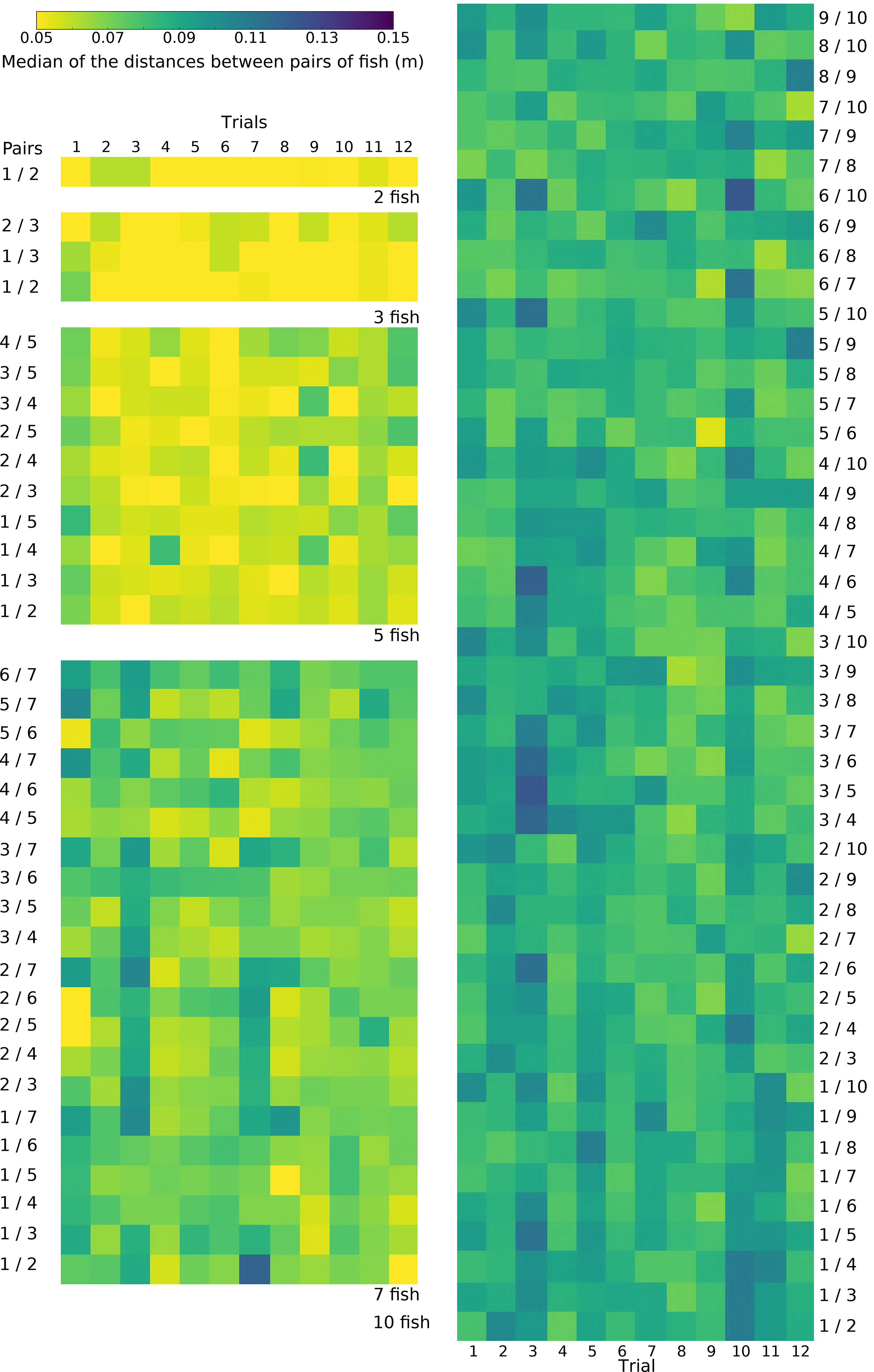

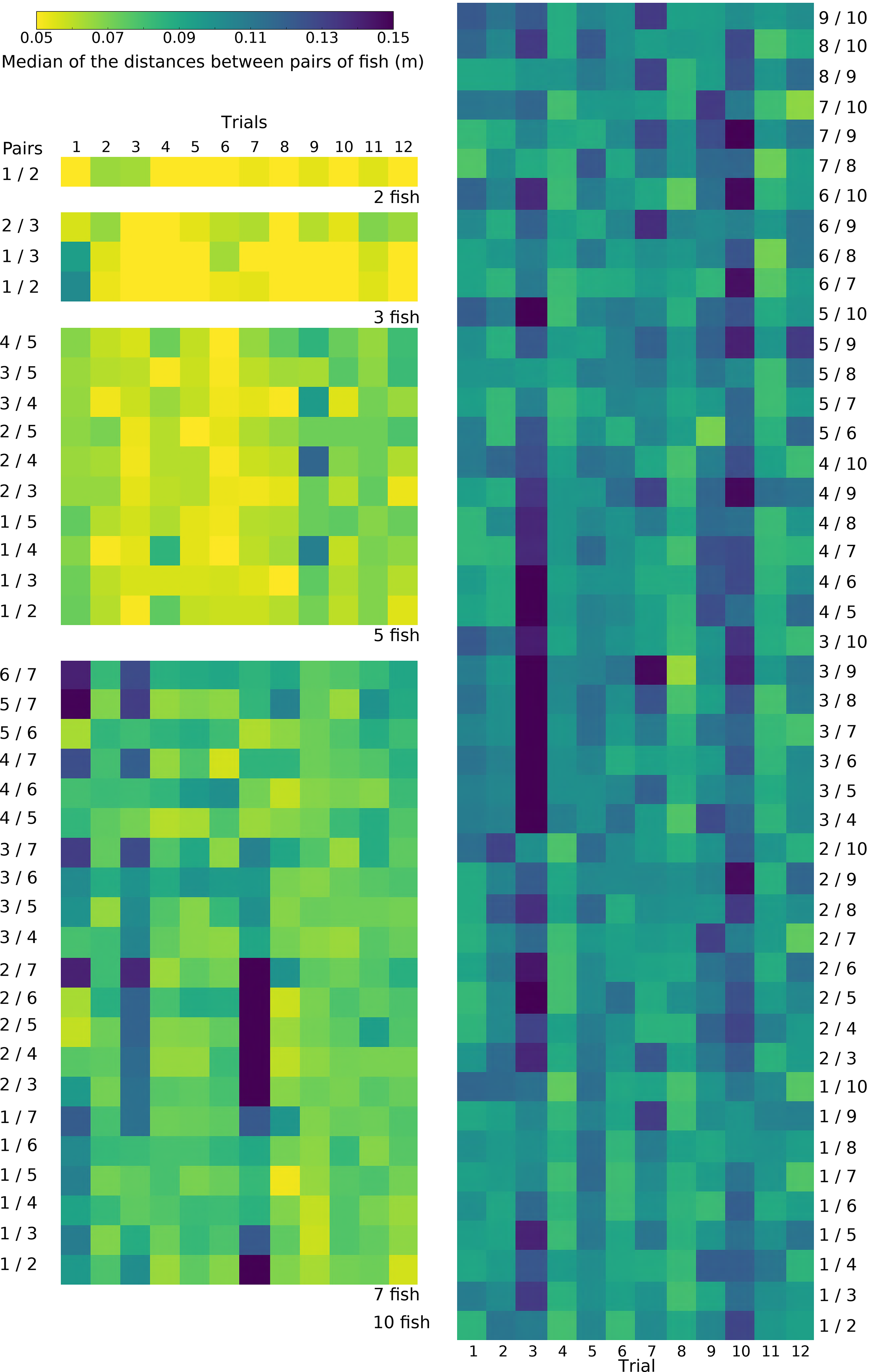

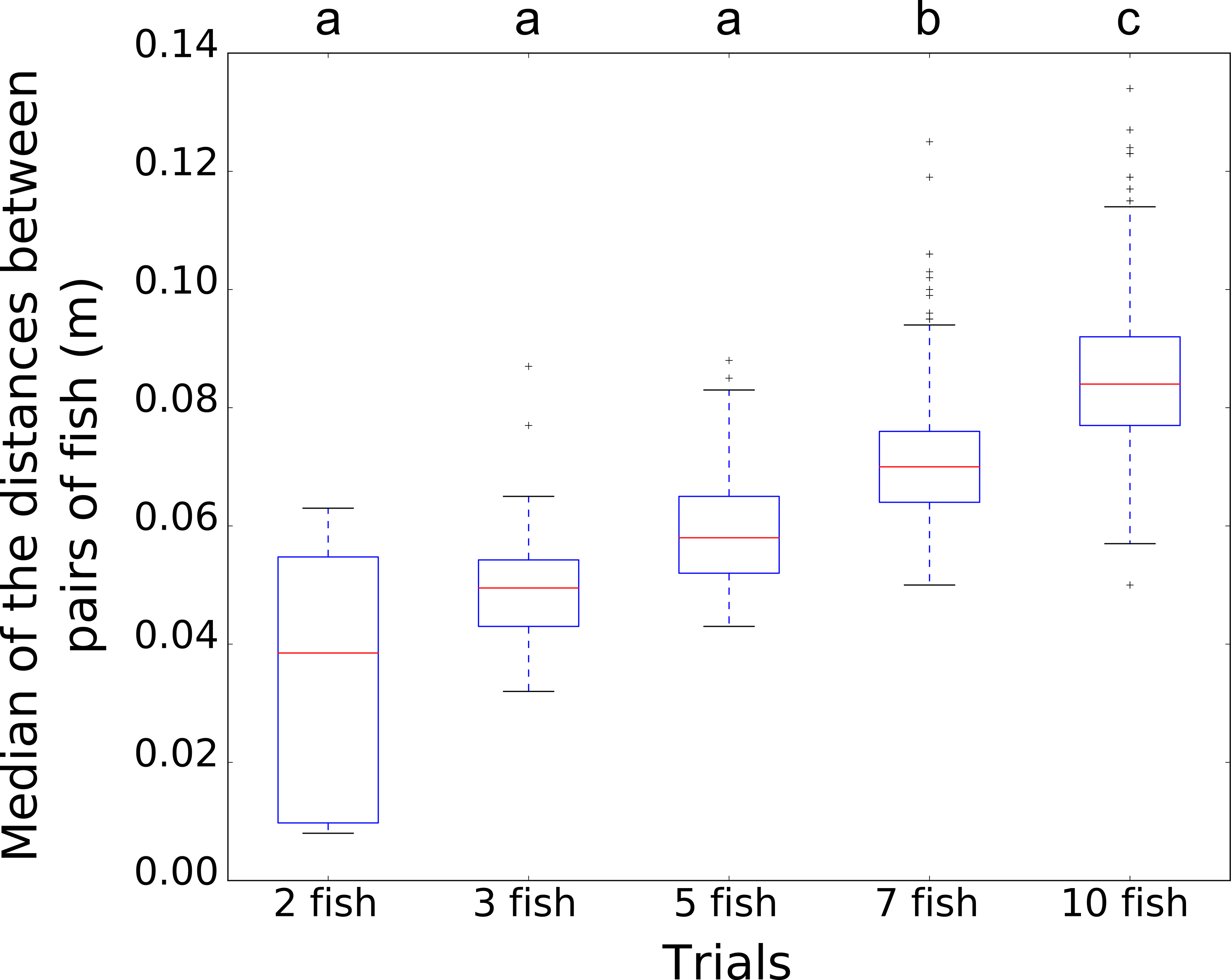

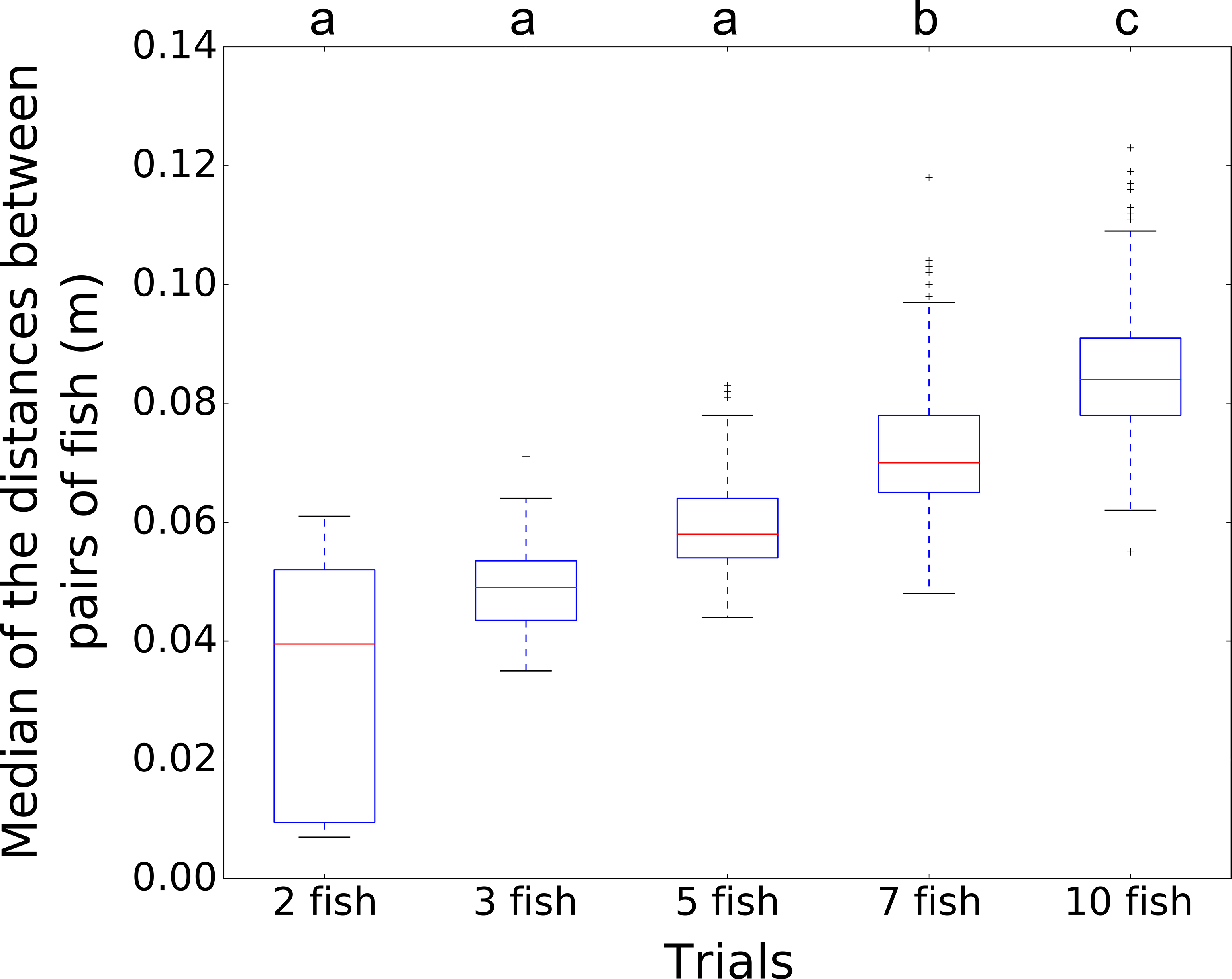

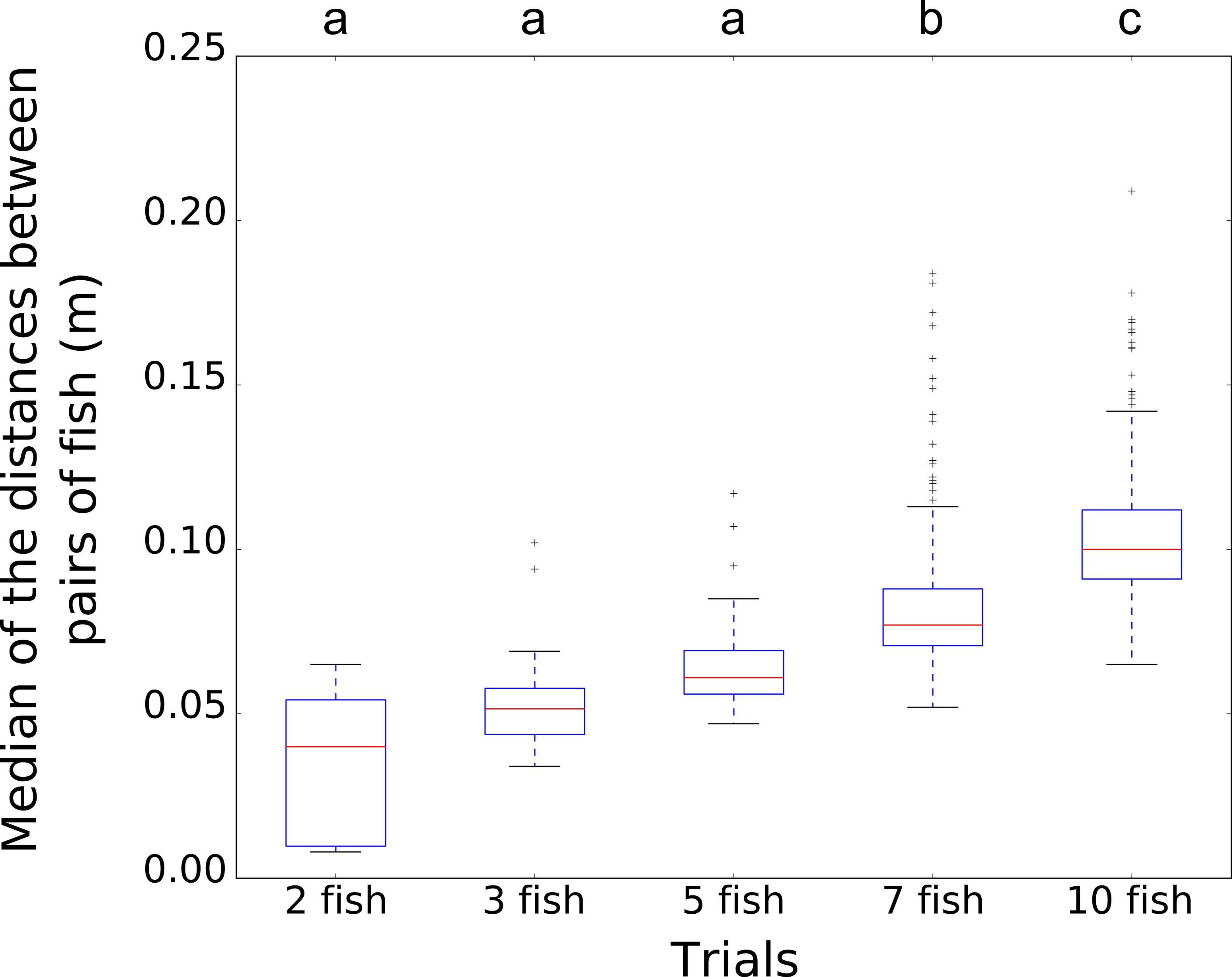

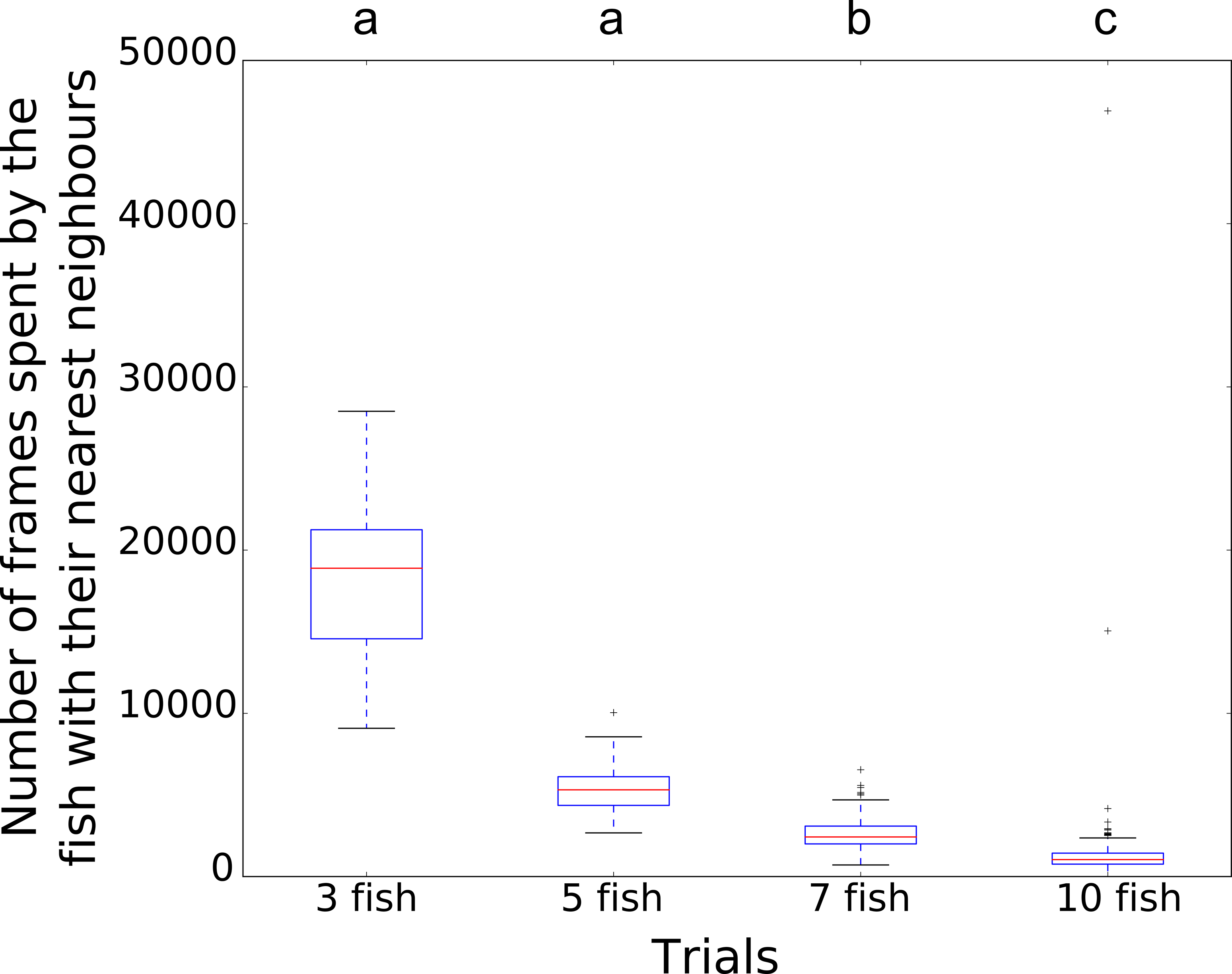

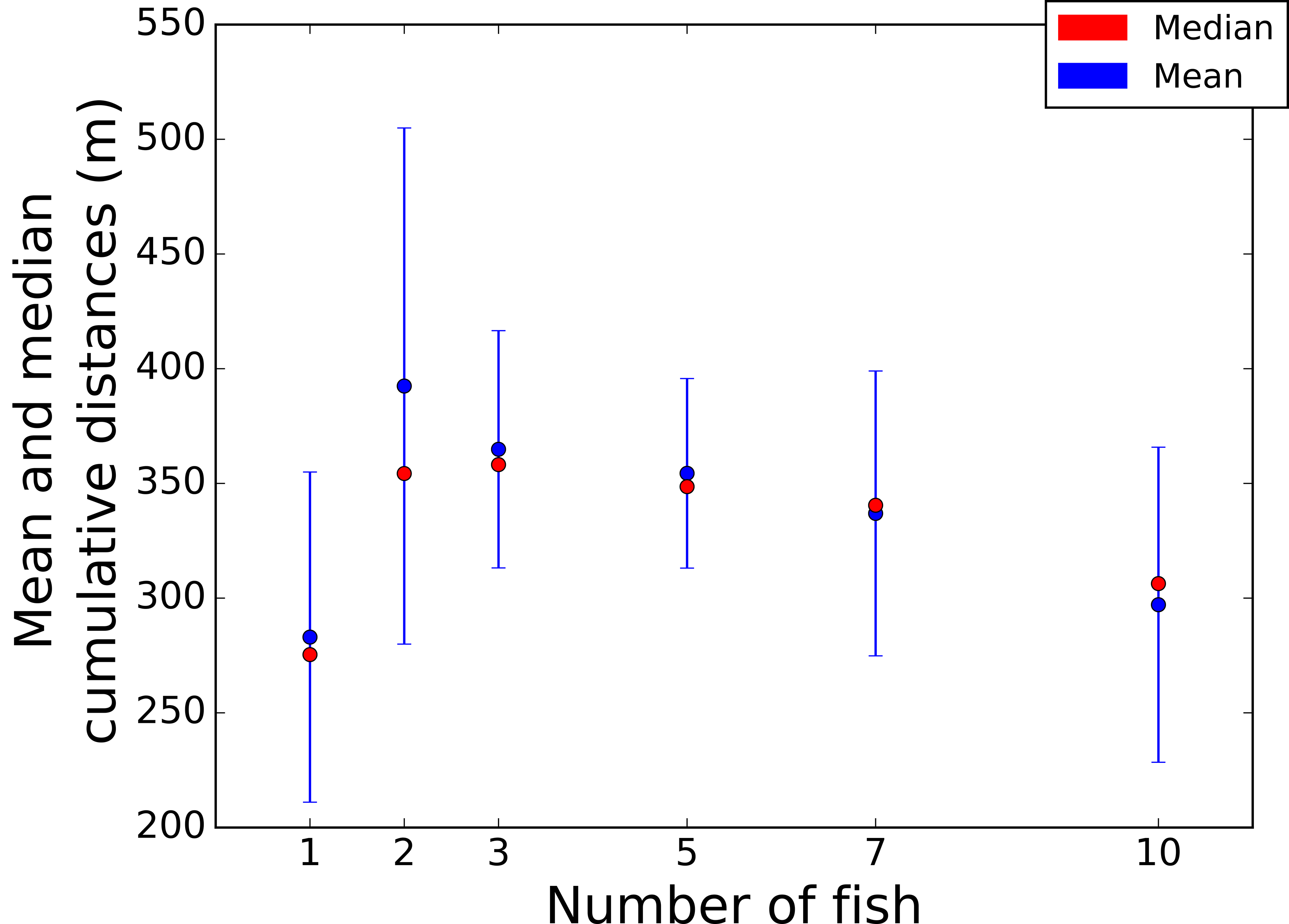

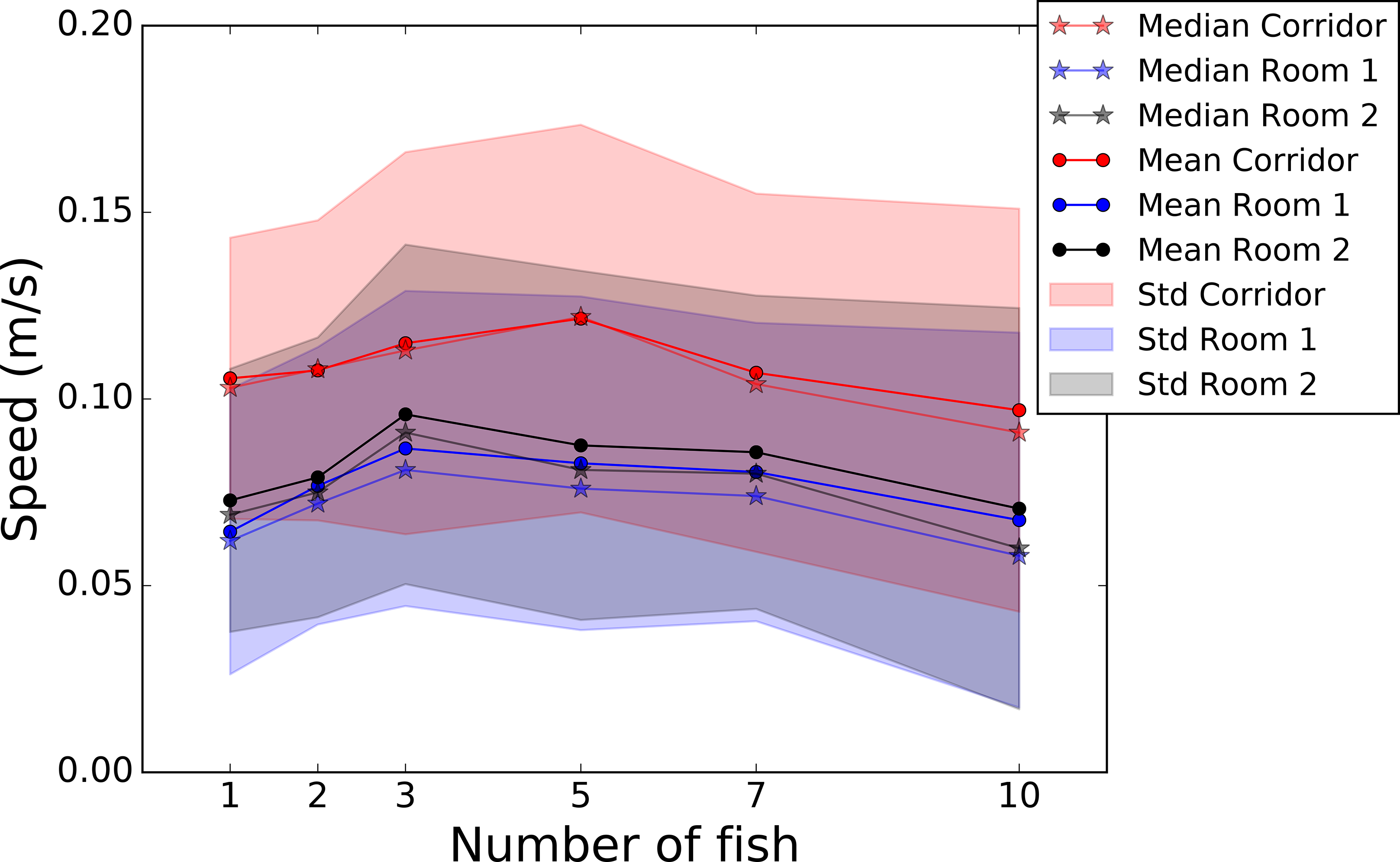

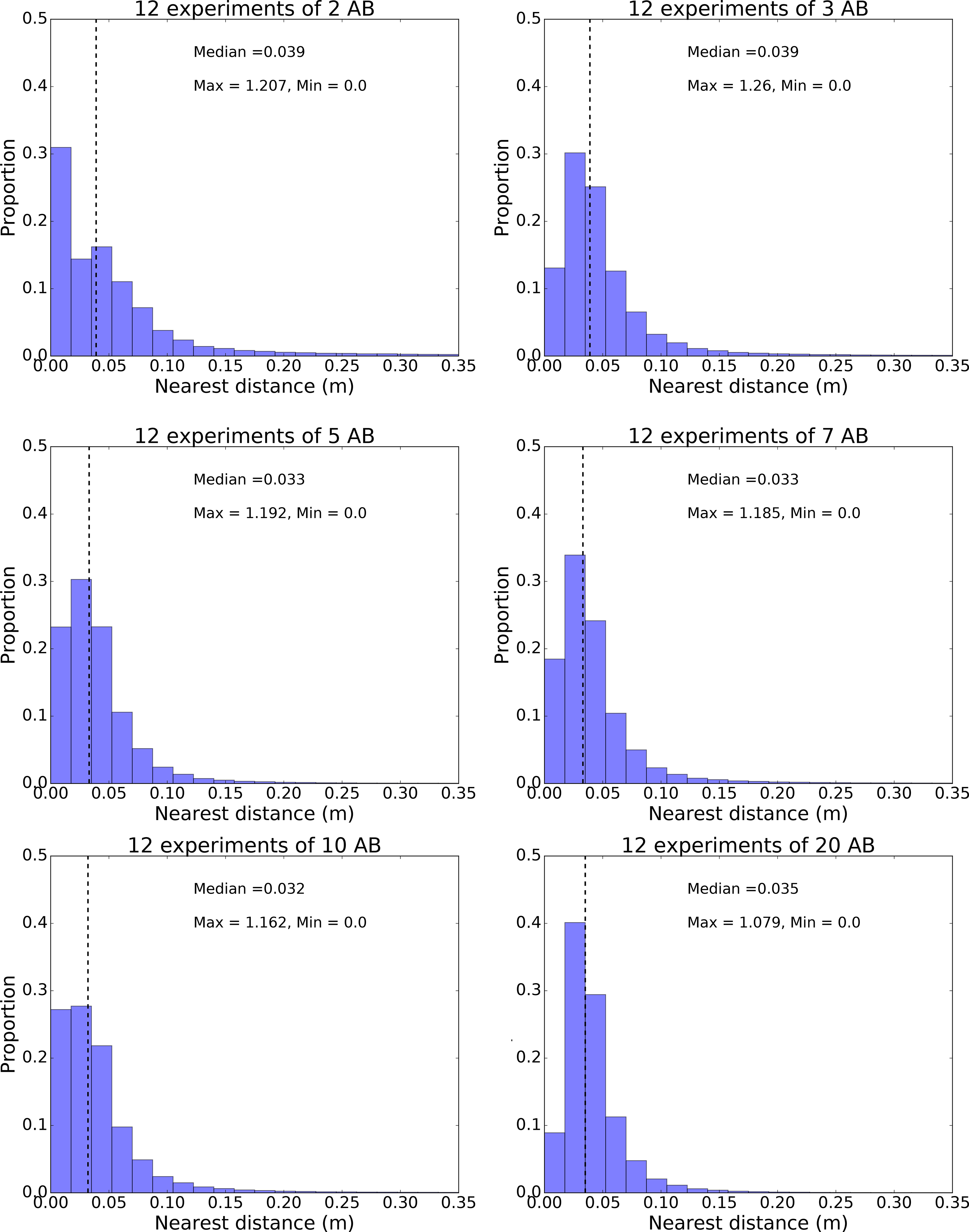

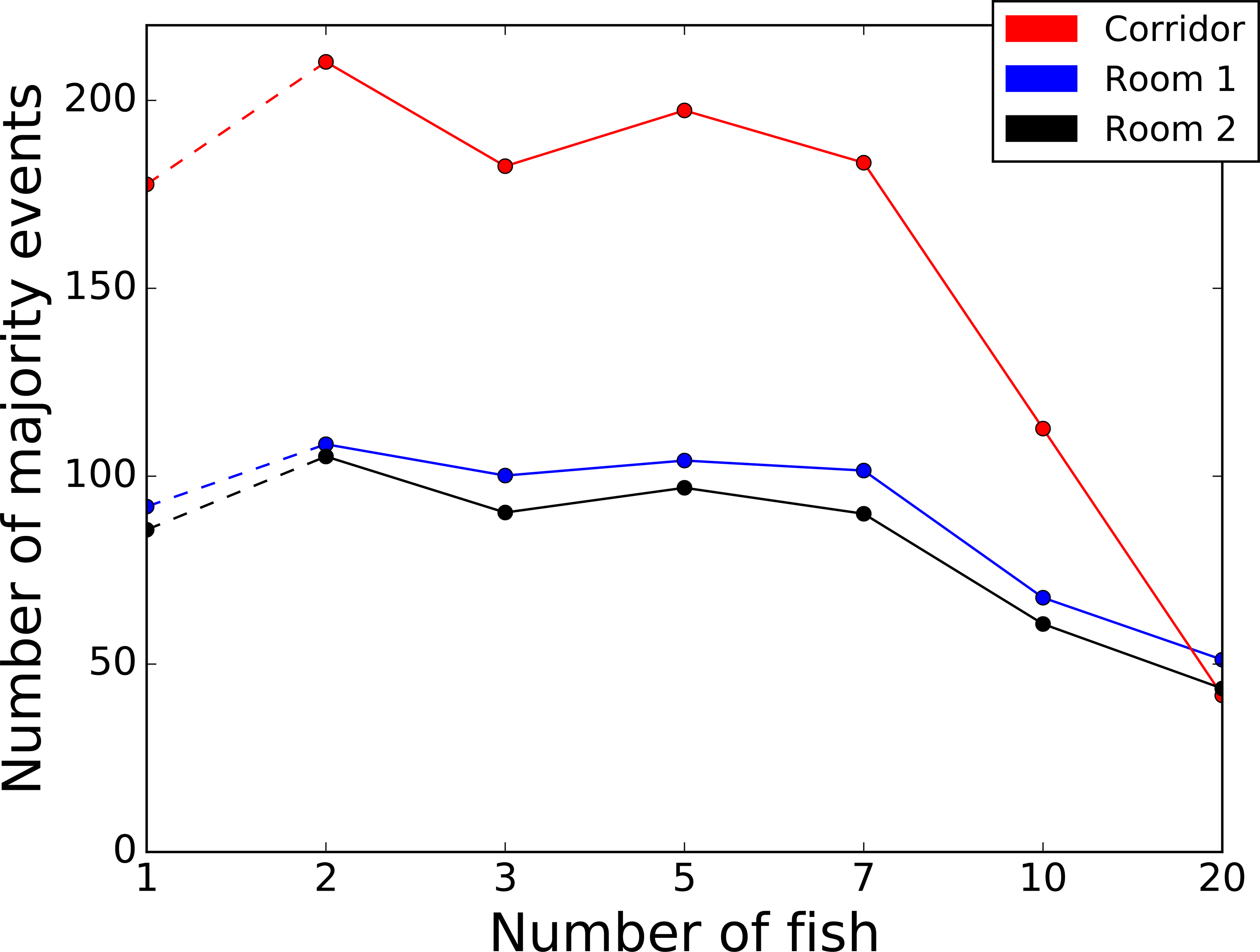

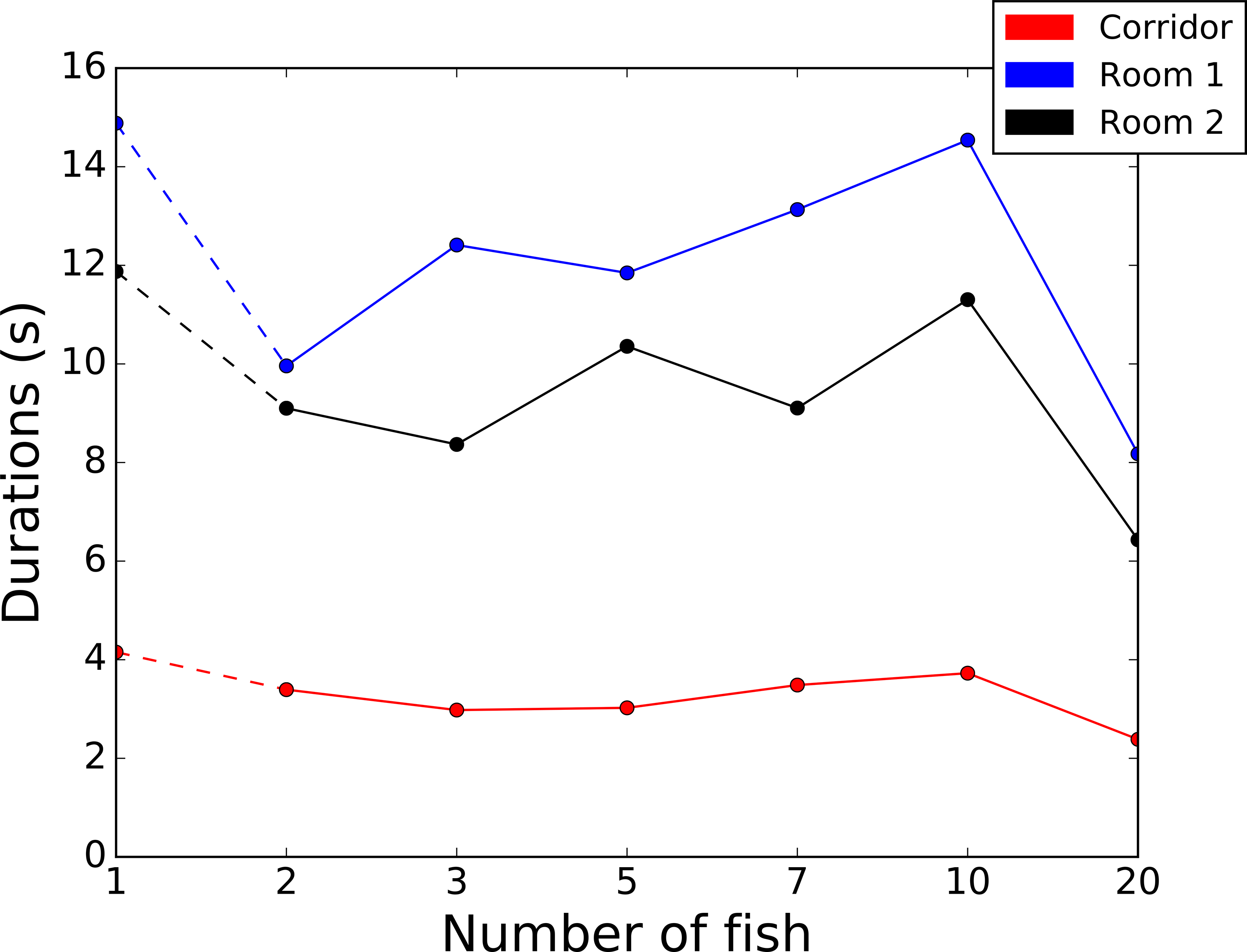

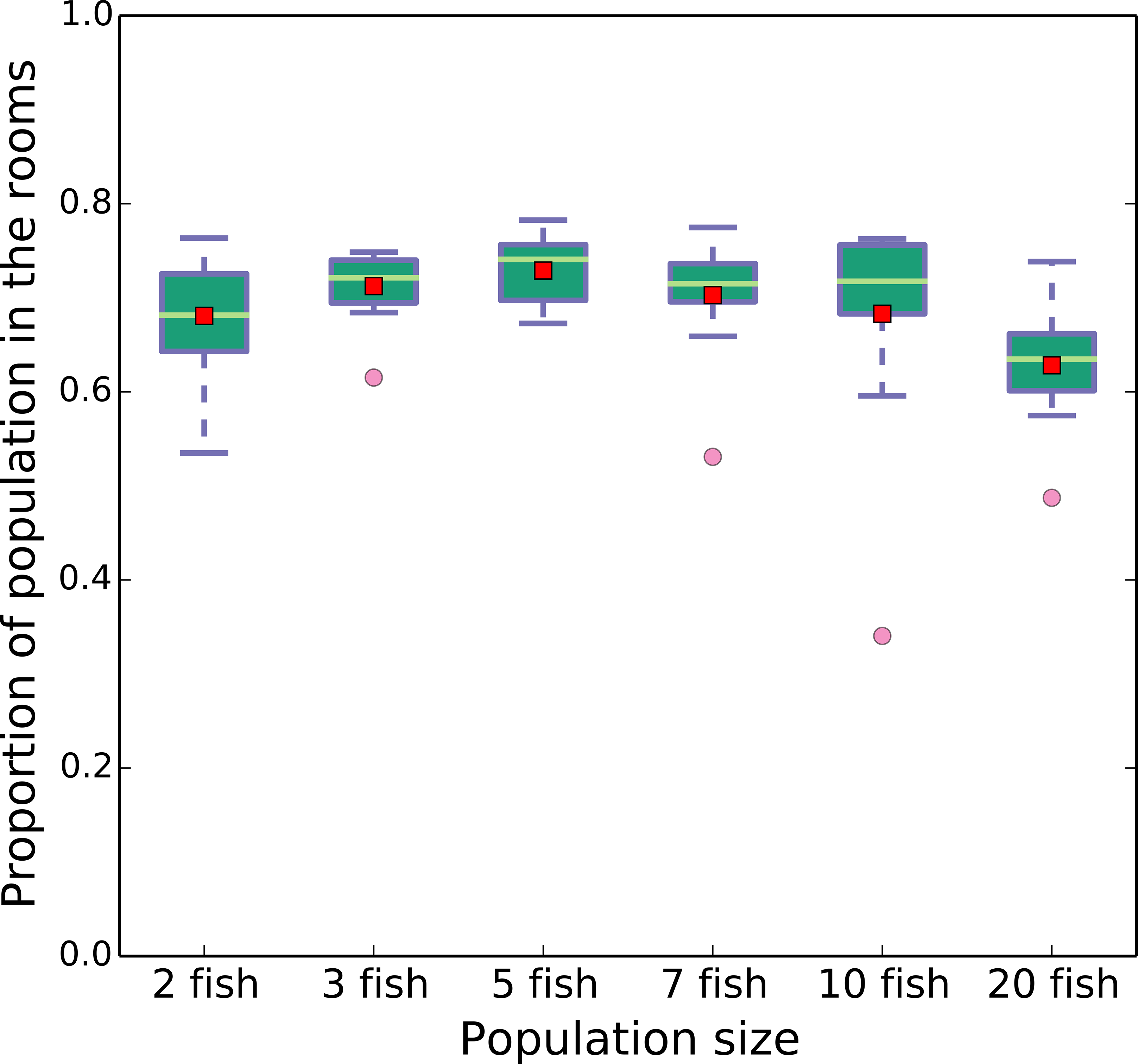

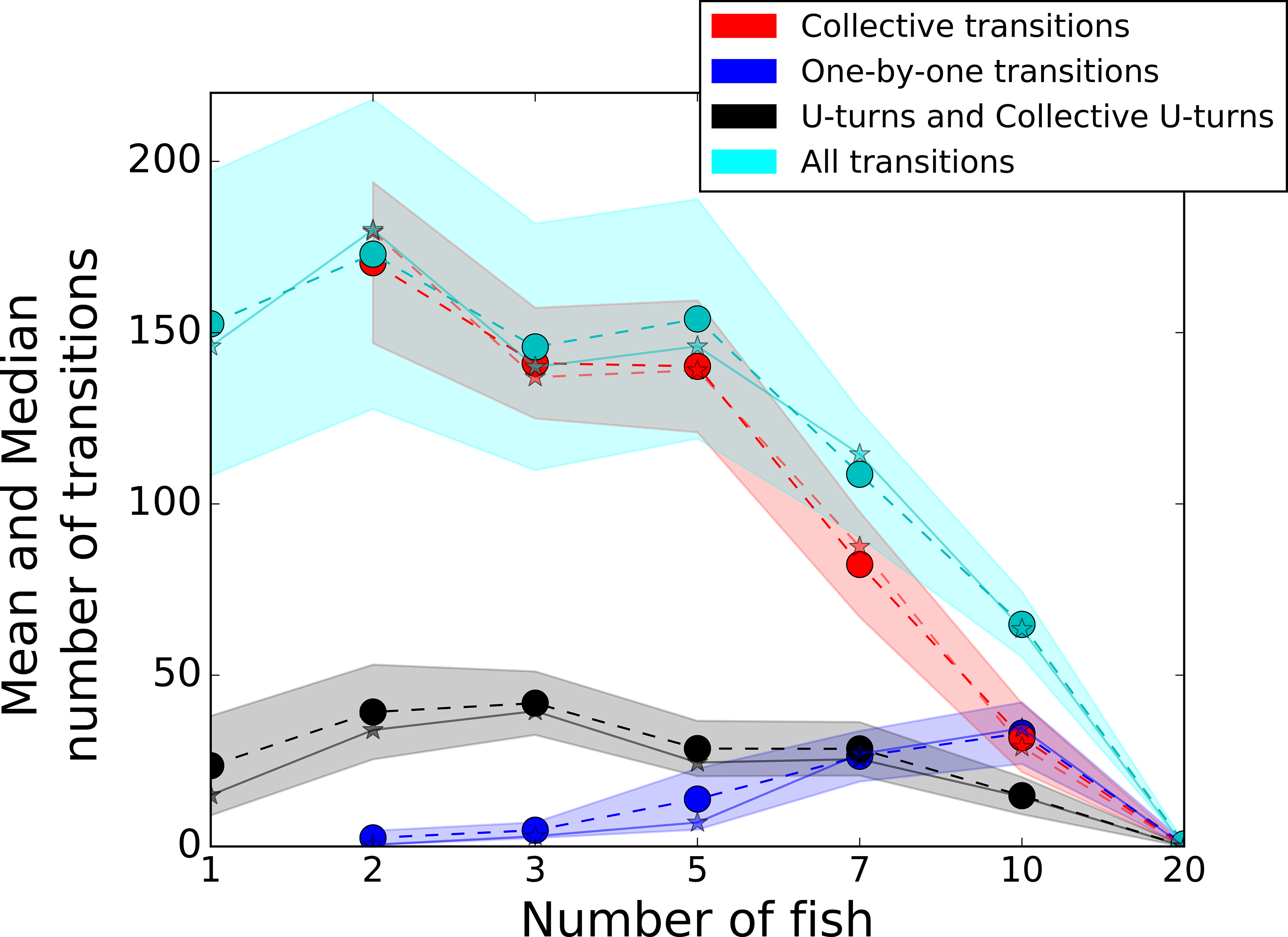

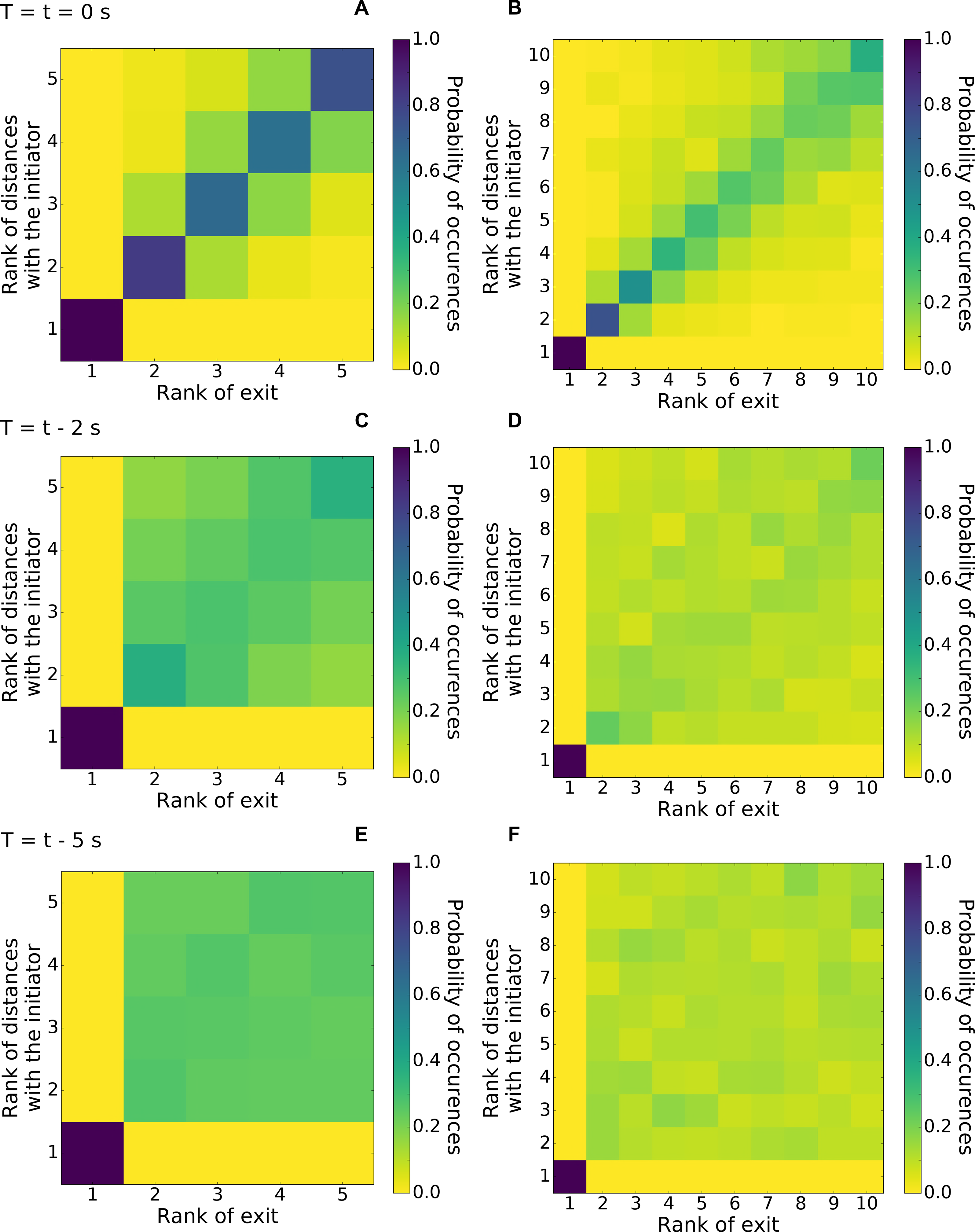

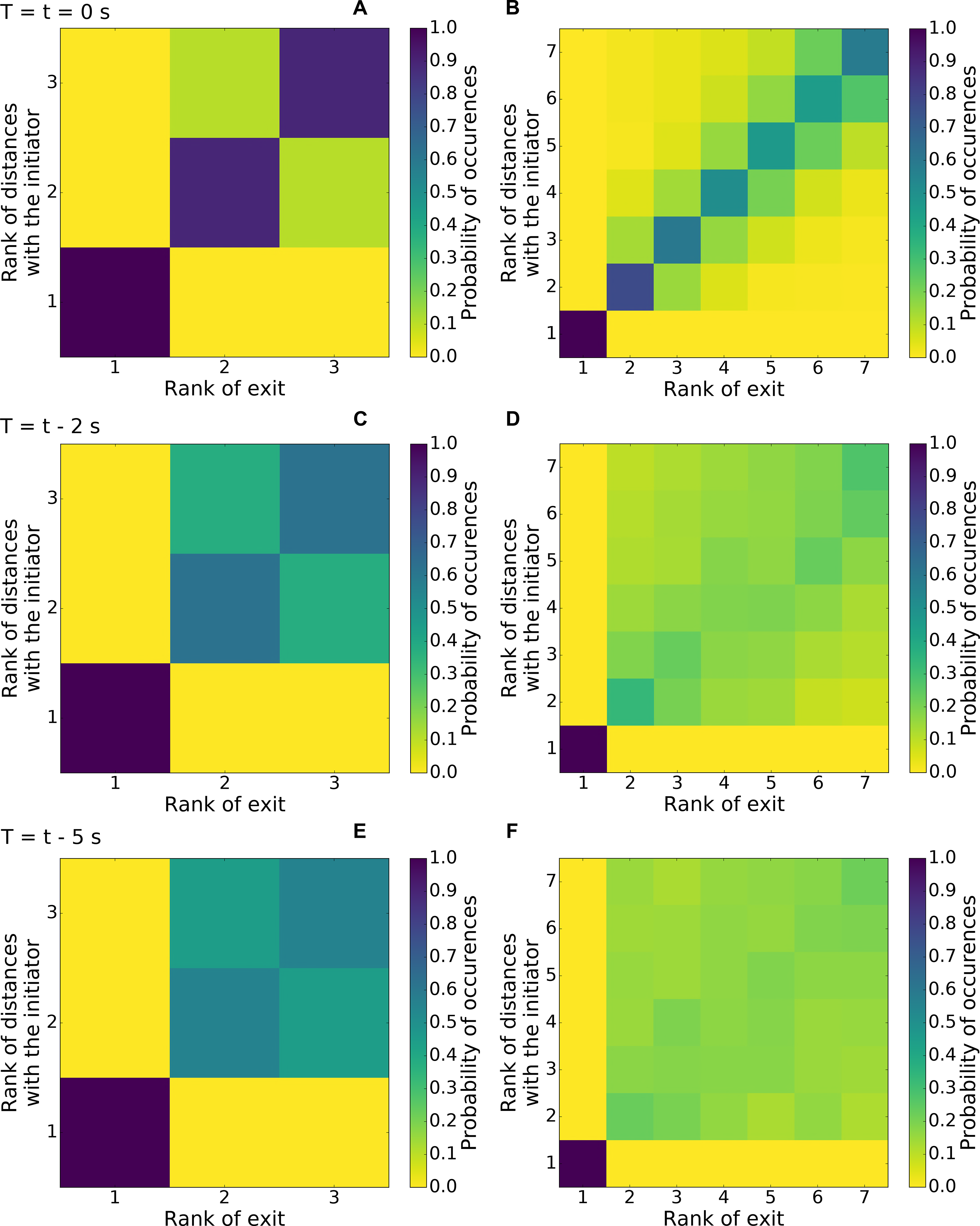

## Bibliography

[1] O. Petit and R. Bon. Decision-making processes: The case of collective movements. Behavioural Processes, 84:635–647, 2010.

[2] C. Sueur, J. L. Deneubourg, and O. Petit. From the first intention movement to the last joiner: macaques combine mimetic rules to optimize their collective decisions. Proceedings of the Royal Society B: Biological Sciences, 278:1697–1704, 2011.

[3] R. E. Engeszer, L. A. Da Barbiano, M. J. Ryan, and D. M. Parichy. Timing and plasticity of shoaling behaviour in the zebrafish, Danio rerio. Animal Behaviour, 74:1269–1275, 2007.

[4] J.K. Parrish, S.V. Viscido, and D. Grübaum. Selforganized fish schools: an examination of emergent properties. Biological Bulletin, 202:296–305, 2002.

[5] J. E. Herbert-Read, A. Perna, R. P. Mann, T. M. Schaerf, D. J. T. Sumpter, and A. J. W. Ward. Inferring the rules of interaction of shoaling fish. Proceedings of the National Academy of Sciences, 108:18726–18731, 2010.

[6] C. K. Hemelrijk and H. Hildenbrandt. Schools of fish and flocks of birds: their shape and internal structure by self-organization. Interface Focus, 2:726–737, 2012.

[7] Dimitrii V. Radakov. Schooling in the ecology of fish. J. Wiley, 1973.

[8] C. Sueur, A. J. King, L. Conradt, G. Kerth, D. Lusseau, C. MettkeHofmann, C. Schaffner, L. Williams, D. Zinner and F. Aureli Collective decision-making and fission-fusion dynamics: a conceptual framework. Oikos, 120(11), 1608–1617, 2011.

[9] N. R. Franks, S. C. Pratt, E. B. Mallon, N. F. Britton, and D. J. T. Sumpter. Information flow, opinion polling and collective intelligence in house-hunting social insects. Philosophical Transactions of the Royal Society of London. Series B, Biological Sciences, 357:1567–1583, 2002.

[10] J.-M. Amé, J. Halloy, C. Rivault, C. Detrain, and J.-L. Deneubourg. Collegial decision making based on social amplification leads to optimal group formation. Proceedings of the National Academy of Science USA, 103:5835–5840, 2006.

[11] G. Rieucau, J. Morand-Ferron and L.-A. Giraldeau. Group size effect in nutmeg mannikin: between-individuals behavioral differences but same plasticity. Behavioural Ecology, 21(4): 684–689, 2010.

[12] S. J. Dahlbom, D. Lagman, K. Lundstedt-Enkel, L. F. Sundström, and S. Winberg. Boldness predicts social status in zebrafish (Danio rerio). PLoS One, 6:e23565, 2011.

[13] J. L. Harcourt, T. Z. Ang, G. Sweetman, R. A. Johnstone and Andrea Manica. Social Feedback and the Emergence of Leaders and Followers. Current Biology, 19:248–252, 2009.

[14] L. Conradt, and T.J. Roper. Consensus decision making in animals. TRENDS in Ecology and Evolution, 20:449–456, 2005.

[15] V. L. Pritchard, J. Lawrence, R. K. Butlin, J. Krause. Shoal choice in zebrafish, Danio rerio: the influence of shoal size and activity Animal Behaviour, 62, 1085–1088, 2001.

[16] R. E. Engeszer, M. J. Ryan and D. M. Parichy. Learned Social Preference in Zebrafish Current Biology, 14:881–884, 2004.

[17] D. J. Hoare, I. D. Couzin, J.-G. J. Godin and J. Krause. Context-dependent group size choice in fish. Animal Behaviour, 67, 155–164, 2004.

[18] I. D. Couzin, J. Krause, N. R. Franks and S. A. Levin Effective leadership and decision-making in animal groups on the move. Nature, 433, 513–516, 2005.

[19] M. Bourjade, B. Thierry, M. Maumy and O. Petit Decision-making in Przewalski horses (Equus ferus przewalskii) is driven by the ecological contexts of collective movements. Ethology, 115:321–330, 2009.

[20] C. Leblond and S. G. Reebs. Individual leadership and boldness in shoals of golden shiners (Notemigonus crysoleucas). Behaviour, 143:1263–1280, 2006.

[21] L. B. Pettersson, C. Brönmark. Trading off safety against food: state dependent habitat choice and foraging in crucian carp. Oecologia, 95:353–357, 1993.

[22] A. J. W. Ward, J. E. Herbert-Read, D. J. T. Sumpter, and J. Krause. Fast and accurate decisions through collective vigilance in fish shoals. Proceedings of the National Academy of Sciences USA, 108:2312–2315, 2011.

[23] C. Kistler, D. Hegglin, H. Wrbel and B. Knig. Preference for structured environment in zebrafish (Danio rerio) and checker barbs (Puntius oligolepis). Applied Animal Behaviour Science, 135:318–327, 2011.

[24] M. Sison and R. Gerlai. Associative learning in zebrafish (Danio rerio) in the plus maze. Behavioural Brain Research, 207:99–104, 2010.

[25] N. Miller, S. Garnier, A. T. Hartnett, and I. D. Couzin. Both information and social cohesion determine collective decisions in animal groups. Proceedings of the National Academy of Sciences USA, 110:5263–5268, 2013.

[26] C. Detrain and J. L. Deneubourg. Collective decisionmaking of foraging patterns in ants and honeybees. Advances in Insect Physiology, 35:123–173, 2008.

[27] A. J. W. Ward, J. E. Herbert-Read, L. A. Jordan, R. James, J. Krause, Q. Ma, D. I. Rubenstein, D. J. T. Sumpter and L. J. Morrell Initiators, Leaders, and Recruitment Mechanisms in the Collective Movements of Damselfish. Am. Nat., 181, 748–760, 2013.

[28] M. Bourjade, B. Thierry, M. Hausberger and O. Petit Is Leadership a Reliable Concept in Animals? An Empirical Study in the Horse. PLoS ONE, 10(5): e0126344, 2015.

[29] B. Collignon, A. Séguret, Y. Chemtob, L. Cazenille, J. Halloy Collective departures in zebrafish: profiling the initiators arXiv preprint, arXiv:1701.03611, 2017.

[30] S. B. Rosenthal, C. R. Twomey, A. T. Hartnett, H. S. Wu and I. D. Couzin Revealing the hidden networks of interaction in mobile animal groups allows prediction of complex behavioral contagion. Proceedings of the National Academy of Sciences, 112(15), 4690–4695, 2015.

[31] S. Nakayama, A. F. Ojanguren and L. A. Fuiman Process-based approach reveals directional effects of environmental factors on movement between habitats. Journal of Animal Ecology, 80, 1299–1304, 2011.

[32] S. Nakayama, J. L. Harcourt, R. A. Johnstone, A. Manica Initiative, Personality and Leadership in Pairs of Foraging Fish. PLoS ONE, 7(5), e36606, 2012.

[33] Ch. Becco, N. Vandewalle, J. Delcourt, P. Poncin Experimental evidences of a structural and dynamical transition in fish school. Physica A, 367:487–493, 2006

[34] J. E. Herbert-Read, S. Krause, L. J. Morrell, T. M. Schaerf, J. Krause and A. J. W. Ward The role of individuality in collective group movement. Proc R Soc B, 280: 2012–2564, 2013.

[35] K. Tunstrom, Y. Katz, C. C. Ioannou, C. Huepe, M. J. Lutz, I. D. Couzin Collective States, Multistability and Transitional Behavior in Schooling Fish. PLOS Computational Biology, 9(2), e1002915, 2013.

[36] D. S. Shelton, B. C. Price, K. M. Ocasio, E. P. Martins Density and group size influence shoal cohesion, but not coordination in zebrafish (Danio rerio). Journal of Comparative Psychology, 129(1), 72, 2015.

[37] J. G. Frommen, M. Hiermes, T. C. M. Bakker Disentangling the effects of group size and density on shoaling decisions of three-spined sticklebacks (Gasterosteus aculeatus). Behavioral Ecology and Sociobiology, 63(8), 11411148, 2009.

[38] A. Séguret, B. Collignon, J. Halloy Strain differences in the collective behaviour of zebrafish (Danio rerio) in heterogeneous environment Royal Society Open Science, 23:R709, 2016.

[39] B. Collignon, A. Séguret, J. Halloy A stochastic vision-based model inspired by zebrafish collective behaviour in heterogeneous environments. Royal Society Open Science, 2016.

[40] R. Jeanson, A. Dussutour, V. Fourcassié Key Factors for the Emergence of Collective Decision in Invertebrates Frontiers in Neuroscience, 6, 121, 2012.

[41] Norton W., Bally-Cuif L. Adult zebrafish as a model organism for behavioural genetics. BMC Neurosci., 11, 90, 2010.

[42] Oliveira R. F. Mind the fish: zebrafish as a model in cognitive social neuroscience. Front. Neural Circuits, 7, 131, 2013.

[43] R. Spence, G. Gerlach, C. Lawrence, and C. Smith. The behaviour and ecology of the zebrafish, Danio rerio. Biological reviews of the Cambridge Philosophical Society, 83:13–34, 2008.

[44] M. M. McClure, P. B. McIntyre, and A. R. McCune. Notes on the natural diet and habitat of eight danionin fishes, including the zebrafish (Danio rerio). Journal of Fish Biology, 69:553–570, 2006.

[45] D.M. Parichy. Advancing biology through a deeper understanding of zebrafish ecology and evolution. eLife, 4:e05635, 2015.

[46] Suriyampola Piyumika S., Shelton Delia S., Shukla Rohitashva, Roy Tamal, Bhat Anuradha, and Martins Emlia P. Zebrafish Social Behavior in the Wild. Zebrafish, 13(1): 1–8, 2016.

[47] R. Spence, M. K. Fatema, M. Reichard, K. A. Huq, M. A. Wahab, Z. F. Ahmed and C. Smith. The distribution and habitat preferences of the zebrafish in Bangladesh. Journal of Fish Biology, 69:1435–1448, 2006.

[48] M. Arunachalam, M. Raja, C. Vijayakumar, P. Malaiammal, R.L. Mayden Natural history of zebrafish (Danio rerio) in India. Zebrafish, 10(1):1–14, 2013.

[49] J. A. Wiens Population responses to patchy environments. Annual review of ecology and systematics, 81–120, 1976.

[50] A. Pérez-Escudero, J. Vicente-Page, R. C. Hinz, S. Arganda, and G. G. de Polavieja. idTracker: tracking individuals in a group by automatic identification of unmarked animals Nature methods, 11(7):743–748, 2014.

[51] C. A. Dlugos, and R. A. Richard Ethanol effects on three strains of zebrafish: model system for genetic investigations Pharmacology Biochemistry and Behavior, 74(2):471–480, 2003.

[52] M. Burd and N. Aranwela Head-on encounter rates and walking speed of foragers in leaf-cutting ant traffic. Insectes soc. 50: 3, 2003.

[53] D. Chowdhury, A. Schadschneider, K. Nishinari Physics of transport and traffic phenomena in biology: from molecular motors and cells to organisms. Physics of Life Reviews Volume 2, Issue 4:318352, 2005.

[54] C. Magnhagen, N. Bunnefeld Express your personality or go along with the group: what determines the behaviour of shoaling perch? Proc. R. Soc. B 276, 33693375, 2009

[55] A. L. J. Burns, J. E. Herbert-Read, L. J. Morrell, A. J. W. Ward Consistency of leadership in shoals of mosquitofish (Gambusia holbrooki) in novel and in familiar environments. PLoS ONE 7, e36567, 2012

[56] S. Nakayama, R. A. Johnstone, A. Manica Temperament and hunger interact to determine the emergence of leaders in pairs of foraging fish. PLoS ONE 7, e43747, 2012.

[57] A. J. King, L. J. Williams, C. Mettke-Hofmann The effects of social conformity on Gouldian finch personality. Animal Behaviour, Volume 99, January 2015, Pages 2531, 2015.

[58] R. J. Egan, C. L. Bergner, P. C. Hart, J. M. Cachat, P. R. Canavello, M. F. Elegante, S. I. Elkhayat, B. K. Bartels, A. K. Tien, D. H. Tien, S. Mohnot, E. Beeson, Glasgow, H. Amri, Z. Zukowska, A. V. Kalueff Understanding behavioral and physiological phenotypes of stress and anxiety in zebrafish. Behavioural Brain Research, Volume 205, Issue 1, 14 December 2009, Pages3844

[59] L. Briard, C. Dorn and O. Petit Personality and affinities play a key role in the organisation of collective movements in a group of domestic horses. Ethology, 121: 888–902, 2015.

[60] C. Sueur and O. Petit Organization of group members at departure is driven by social structure in Macaca. Int. J. Primatol., 29, 1085–1098, 2008.

[61] C. Sueur, O. Petit and J. Deneubourg Selective mimetism at departure in collective movements of Macaca tonkeana: an experimental and theoretical approach. Anim. Behav., 78, 1087–1095, 2015.

[62] A. J. King, D. D. P. Johnson and M. Van Vugt A rule-of-thumb based on social affiliation explains collective movements in desert baboons. Anim. Behav., 82, 1337–1345, 2011.

[63] J. W. Jolles, N. J. Boogert, V. H. Sridhar, I. D. Couzin, A. Manica Consistent Individual Differences Drive Collective Behavior and Group Functioning of Schooling Fish Current Biology, 27 (18), 28622868.e7, 2017.

[64] D. J. Hoare, J. Krause, N. Peuhkuri and J.-G. J Godin Body size and shoaling in fish. Journal of Fish Biology 57, 1351–1366, 2000

[65] J. Krause and G. D. Ruxton Living in groups. Oxford Univ. Press, 2002.

[66] D. P. Croft, B. J. Arrowsmith, J. Bielby, K. Skinner, E. White, I. D. Couzin, A. E. Magurran, I. Ramnarine and J. Krause Mechanisms underlying shoal composition in the Trinidadian guppy, Poecilia reticulata. Oikos, 100(3), 429–438, 2003.

[67] M. H. Pillot, J. Gautrais, J. Gouello, P. Michelena, and R. Bon Moving together: Incidental leaders and nave follower. Behavioural Processes, 83(3), 235–241, 2010.

[68] R. W. Byrne, A. Whiten, S. P. Henzi Social relationships of mountain baboons: leadership and affiliation in a nonfemale-bonded monkey. Am. J. Primatol., 20, 313–329, 1990.

[69] J.-B. Leca, N. Gunst, N. Thierry, O. Petit Distributed leadership in semifree-ranging white-faced capuchin monkeys. Animal Behaviour, 66(6), 1045–1052, 2003.

[70] O. Petit, J. Gautrais, J.-B. Leca, G. Theraulaz, and J.-L. Deneubourg. Collective decision-making in white-faced capuchin monkeys. Proceedings of the Royal Society of London B: Biological Sciences, 276(1672):3495–3503, 2009.

[71] A. J. W. Ward, D. J. T. Sumpter, I. D. Couzin, P. J. B. Hart, and J. Krause Quorum decision-making facilitates information transfer in fish shoals. PNAS, 105 (19), 6948–6953, 2008.

[72] K. Branson, A. A Robie, B. Bender, P. Perona, and H. M. Dickinson. High-throughput ethomics in large groups of drosophila. Nature, 6:451–457, 2009.

[73] H. Dankert, L. Wang, E. D. Hoopfer, D. J. Anderson, and P. Perona. Automated monitoring and analysis of social behavior in drosophila. Nature methods, 6:297–303, 2009.

[74] M. Ballerini, N. Cabibbo, R. Candelier, A. Cavagna, E. Cisbani, I. Giardina, V. Lecomte, A. Orlandi, G. Parisi, A. Procaccini, M. Viale, and V. Zdravkovic. Interaction ruling animal collective behavior depends on topological rather than metric distance: Evidence from a field study. Proceedings of the National Academy of Sciences USA 105:1232–1237, 2007.

[75] N. Miller and R. Gerlai. Quantification of shoaling behaviour in zebrafish (Danio rerio). Behavioural brain research, 184:157–166, 2007.

[76] N. Miller and R. Gerlai. Shoaling in zebrafish: what we don’t know. Reviews in the neurosciences, 22:17–25, 2011.

[77] A. Strandburg-Peshkin, C.R. Twomey, N.W.F. Bode, A. B. Kao, Y. Katz, C.C Ioannou, S.B. Rosenthal, C.J. Torney, H.S. Wu, S.A. Levin, and I.D. Couzin. Visual sensory networks and effective information transfer in animal groups. Current Biology, 23:R709, 2013.

[78] M. G. Kendall. A New Measure of Rank Correlation. Biometrika, 30:81–93, 1938.

